# Recall of B cell memory depends on relative locations of prime and boost

**DOI:** 10.1101/2021.12.17.472609

**Authors:** Masayuki Kuraoka, Chen-Hao Yeh, Goran Bajic, Ryutaro Kotaki, Shengli Song, Stephen C. Harrison, Garnett Kelsoe

## Abstract

Re-entry of memory B cells to recall germinal centers (GCs) is essential for updating their B-cell antigen receptors (BCRs). Using single B-cell culture and fate-mapping, we have characterized BCR repertoires in recall GCs following boost immunizations at sites local or distal to the priming. Local boosts with homologous antigen recruit to recall GCs progeny of primary GC B cells more efficiently than do distal boosts. Recall GCs following local boosts contain significantly more B cells with elevated levels of *Ig* mutations and higher avidity BCRs. This local preference is unaffected by blockade of CD40:CD154 interaction that terminate active, primary GC responses. Local boosts with heterologous antigens elicit secondary GCs with B-cell populations enriched for cross-reactivity to the priming and boosting antigens; in contrast, cross-reactive GC B cells are rare following distal boosts. Our findings indicate the importance of locality in humoral immunity and inform serial vaccination strategies for evolving viruses.

**One Sentence Summary:** The participation of memory B cells in recall germinal centers depends on whether the boost is local or distal to the priming site.

## INTRODUCTION

Vaccinations or microbial infections activate the adaptive immune system and establish, predominantly through germinal center (GC) responses, long-lasting protective immunity. This durable immunity comprises long-lived plasma cells and memory B (Bmem) cells. The former maintain circulating antibody (Ab) to provide a first line of defense against reinfection. The latter exhibit multiple fates on re-encountering antigen; Bmem cells either proliferate and differentiate into short-lived plasmablasts/-cytes (PBs/PCs) or (re)enter GCs in which they undergo new rounds of antigen-driven selection and *Ig* somatic hypermutation (SHM) to “update” their B-cell receptors (BCRs). Rapid Bmem differentiation into PBs/PCs contributes to prompt, high affinity Ab responses and protective activity, the Bmem cells that re-enter GCs and emerge with updated BCRs play critical roles in combatting evolving viruses (*e*.*g*., HIV-1, influenza, SARS-CoV-2). Viruses that accumulate fitness-enhancing mutations in response to immune pressure (in individuals or populations) can generate escape variants to which pre-existing Bmem BCRs and circulating Abs bind weakly or not at all.

Studies in mice and humans have provided substantial evidence for the recruitment of Bmem cells into recall GCs (*1-4*) and for the updating of Bmem BCRs (*3, 5-7*). Nonetheless, questions have arisen concerning participation of Bmem cells in recall GC responses (*8-10*), and recent studies have suggested that recruitment of Bmem cells to secondary GCs is not an efficient process. (*11*). Fate-mapping experiments have shown that the marked, progeny of primary GC B cells account for a small subset (≤5%) of secondary GC B cells and that newly-activated, mature B cells dominate the recall GC responses (*11, 12*). In addition to this overall inefficiency, recruitment of high affinity Bmem cells into recall GCs may be further reduced as higher affinity Bmem cells appear to be biased for PBs/PCs differentiation rather than GC re-entry (*13-18*).

That immunization or microbial infection establishes humoral memory that is local as well as systemic is now recognized. For example, pulmonary infection of mice with influenza viruses establishes resident Bmem cells in the lung, and these local Bmem cells exhibit distinct BCR specificities and contribute to early local plasmacytic responses that provide stronger protection against influenza challenge than do systemic Bmem cells (*19, 20*). This property of local humoral memory may be general. Early studies report that antigen-retention in local LNs has significant roles in the migration and retention of specific Bmem cells and that LNs linked to primary immunization sites produce more Ab-secreting cells than do distal LNs after secondary challenge (*21-24*).

Does a distinct population of local Bmem cells in peripheral LNs support secondary GC responses that are distinct from those at distal sites? Is the importance of location in humoral immunity relevant only to non-lymphoid tissues, *e*.*g*., lung, or does it extend to secondary lymphoid tissues? To address these questions, we compared secondary GC responses to boost immunizations given in the same (ipsilateral) or opposite (contralateral) leg of primed mice. The magnitude of secondary GC and serum Ab responses to homologous boosts were similar for both ipsilateral and contralateral boosts. We found, however, that the quality of these responses differed. Ipsilateral boosts elicited GCs with higher numbers of cells that were the progeny of the primary GC response than did contralateral boosts. Consequently, secondary GCs elicited by ipsilateral boosting contained higher numbers of B cells with elevated *Ig* mutation frequencies and higher avidity for antigen. Disruption of active, primary GC responses by injecting anti-CD154 Ab did not reduce this local recall bias for previously mutated, higher affinity cells. In response to boosts with heterologous antigens, ipsilateral boosts were more effective in producing secondary GC B cells that bound both the primary and boost antigens than did contralateral boosts. Our findings suggest that local B-cell memory is retained in the form of GC B cells and/or GC-phenotype B cells. The results have implications for strategies of vaccination against evolving pathogens, for which updating BCRs is essential for durable protection.

## RESULTS

### Boost immunizations at local sites and at distal sites elicit comparable levels of recall serum IgG

To compare Ab responses following boost immunizations at local and distal sites, we immunized B6 mice with influenza hemagglutinin (HA) H1 SI-06 in the right footpad, and then boosted these animals 1-3 month(s) later with homologous HAs either in the right hock (ipsilateral boosts) or in the left hock (contralateral boosts). Eight days after boosting, we quantified HA-specific IgG Abs in sera by a Luminex multiplex bead assay, and enumerated PBs/PCs in the draining LNs by flow cytometry (Fig. 1A). Boost immunizations raised the concentrations of HA-specific serum IgG Abs by ∼12-fold (*p* < 0.001; ipsilateral boosts, 14-fold; contralateral boosts, 10-fold; Figs. 1B and 1C), but there were no significant differences in H1 HA-specific serum IgGs between ipsilateral and contralateral boosts (*p* > 0.99; Figs. 1B and 1C).

**Fig. 1.**
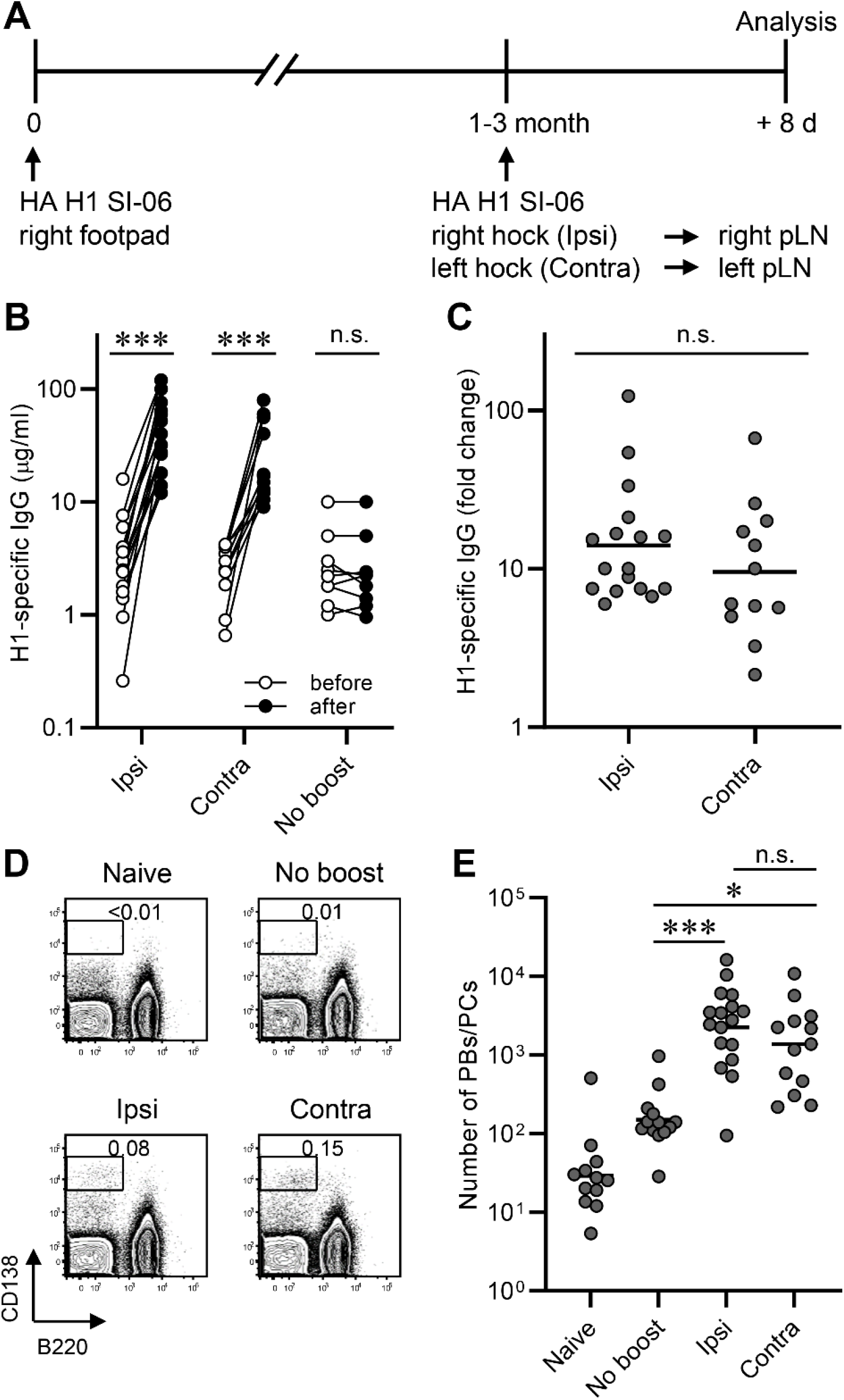
Boost immunizations at local distal sites elicit comparable levels of recall serum IgG. Plasmacytic responses following boosts with homologous antigens. (**A**) Graphic representation of the immunization strategy used in experiments in Figs. 1-4. (**B**) Concentrations of serum IgGs specific to H1 SI-06 before and after boosts ipsilaterally (n = 17) or contralaterally (n = 12). No-boost group (n = 9) received primary immunizations but not boosts. Each symbol represents an individual mouse, and serum samples from the same animals are connected with lines. ***, *p* < 0.001; n.s., *p* > 0.05 by Wilcoxon matched-pairs signed rank test. (**C**) Fold changes in concentrations of H1 HA-specific IgGs in serum samples in (**B**) after boosts. n.s., *p* > 0.05 by Mann-Whitney’s U test. (**D** and **E**) Representative flow diagrams (**D**) and number (**E**) of B220^lo^CD138^hi^ PBs/PCs in the draining LNs from naïve (n = 12), No-boost (n = 13), and boosted mice (ipsilateral boost, n = 17; contralateral boost, n = 13). Numbers near boxes in (**D**) represent frequency of B220^lo^CD138^hi^ cells among live lymphocyte population. ***, *p* < 0.001; *, *p* < 0.05; n.s., *p* > 0.05 by Kruskal-Walis test with Dunn’s multiple comparisons. Combined data from 9 independent experiments are shown.

Consistent with robust serum IgG responses, the number of B220^lo^CD138^hi^ PBs/PCs in the draining LNs were higher after boosting, ∼15-fold (*p* = 0.001) and ∼8-fold (*p* = 0.038) following ipsilateral and contralateral boosts, respectively than in no-boost controls (Figs. 1D and 1E). Between boost regimens there were no significant differences in the number of PBs/PCs in the draining LNs (*p* > 0.99; Fig. 1E). We sorted these samples into three groups by intervals between the priming and boosting (*i*.*e*., 4-5 week, 8-10 week, and 12-14 week intervals, respectively) and found across all intervals that ipsilateral boosts and contralateral boosts elicited comparably robust serum IgG responses and plasmacytic responses in the draining LNs (Fig. S1).

### Secondary GCs following local boosts contain high avidity, antigen-specific B cells at elevated frequency

We characterized secondary GC responses in the draining LNs following ipsilateral or contralateral boosts by enumerating the number of B220^+^CD138^-^GL-7^+^CD38^lo^IgD^-^ GC B cells and by determining their BCR reactivity and avidity. Flow cytometric analysis showed that GC responses elicited by the primary immunization had waned largely, but not completely, by the time of the boosts (Fig. 2A and 2B). We recovered ∼6 times as many GC phenotype B cells from the draining LNs of no-boost controls (geometric mean = 1,700) than from naïve controls (geometric mean = 270). Although this difference was not statistically significant (*p* = 0.72), ∼85% (11 out of 13) of LN samples from no-boost controls contained more than 1,000 GC B cells, while only one third (4 out of 12) of LN samples from naïve mice did (Fig. 2B).

**Figure 2.**
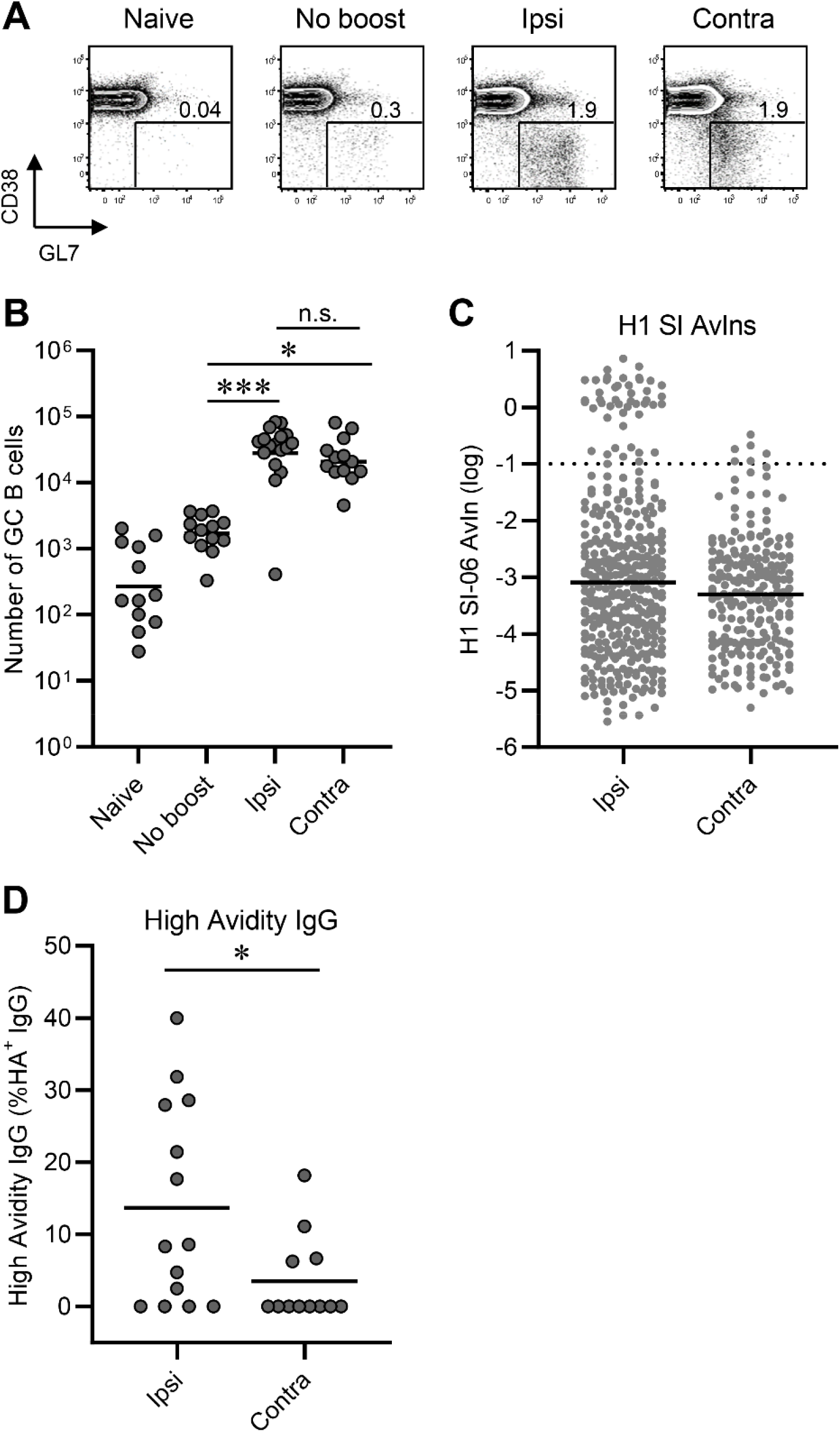
Secondary GCs following local boosts contain a greater proportion of high avidity, antigen-specific B cells than do those following distal boosts. GC responses following boosts with homologous antigens (see also a legend of Fig. 1A). (**A** and **B**) Representative flow diagrams of GL-7 and CD38 expressions on B220^+^CD138^-^ cells (**A**) and number of B220^+^CD138^-^GL-7^+^CD38^lo^IgD^-^ GC B cells in the draining LNs of naïve (n = 12), No-boost (n = 13), and boosted mice (ipsilateral boosts, n = 17; contralateral boosts, n = 13). Numbers near boxes in (**A**) represent frequency of GL-7^+^CD38^lo^IgD^-^ cells among B220^hi^CD138^-/lo^ cells. Horizontal bars in (**B**) represent geometric mean. ***, *p* < 0.001; *, *p* < 0.05; n.s., *p* > 0.05 by Kruskal-Walis test with Dunn’s multiple comparisons. Combined data from 9 independent experiments are shown. (**C**) Distributions of AvIn values for H1 HA-specific IgGs from single-cell cultures of GC B cells are shown (n = 368 and 214 for ipsilateral and contralateral boosts, respectively). Each dot represents individual culture supernatant IgG. Horizontal bars represent geometric mean. Dotted line represents AvIn = 0.1, which we considered high avidity. Combined data from 6-8 independent experiments are shown. (**D**) Frequency of high avidity IgGs (AvIn > 0.1) among HA H1 SI-06 specific IgGs. Each dot represents individual mouse (n = 14, and 12 for ipsilateral and contralateral boosts, respectively). *, *p* < 0.05 by Mann-Whitney’s U test.

Both ipsilateral and contralateral boosts elicited strong secondary GC responses; numbers of GC B cells were ∼17-fold (*p* < 0.001) and ∼13-fold (*p* = 0.011) higher following ipsilateral and contralateral boosts, respectively, than in no-boost controls (Figs. 2A and 2B). Ipsilateral and contralateral boosts elicited secondary GC responses of comparable magnitudes (*p* > 0.99; Fig. 2A and 2B). This relationship held when we boosted mice at 4-5 weeks, 8-10 weeks or 12-14 weeks after priming (Fig. S2A-S2C).

To determine the specificity and affinity of B cells in secondary GCs, we sorted individual GC B cells from the draining LNs of boosted animals into single-cell Nojima cultures (Kuraoka et al., 2016). After culture, we screened clonal IgG Abs in culture supernatants by a Luminex multiplex assay for binding to HA H1 SI-06 (Fig. 2C). We obtained 5,113 clonal IgG examples from secondary GC B cells following ipsilateral (n = 2,255) or contralateral boosts (n = 2,858) (Table 1). In contrast to the similar magnitudes of GC responses induced by ipsilateral or contralateral boosting, the frequency of antigen-specific clonal IgGs from secondary GC B cells elicited by ipsilateral boosts was significantly (*p* = 0.0011) higher [16% (±5.9%)] than that of GC B cells following contralateral boosts ([7.7% (±4.7%]); Fig. S2D, Table 1).

**Table 1.**
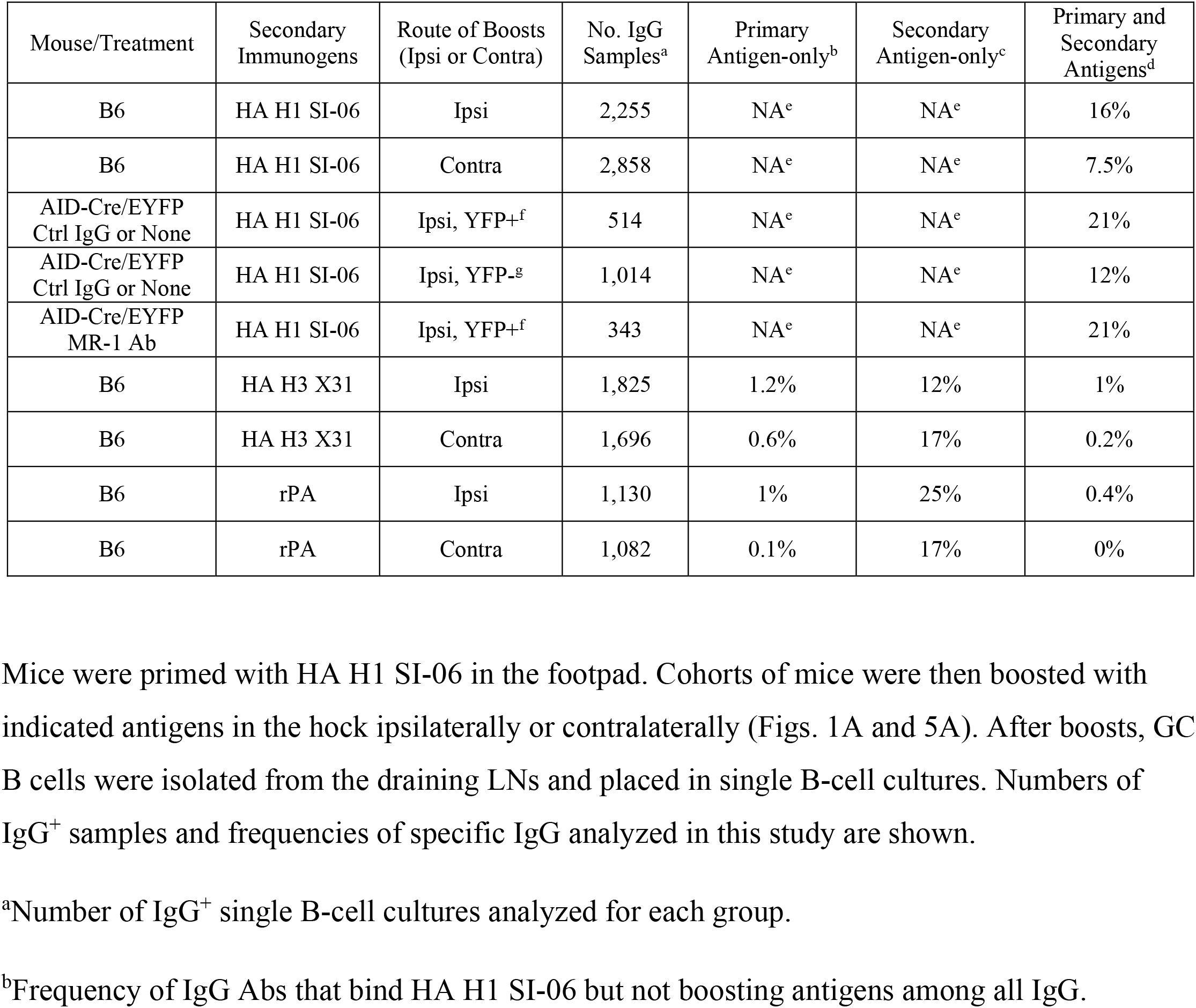
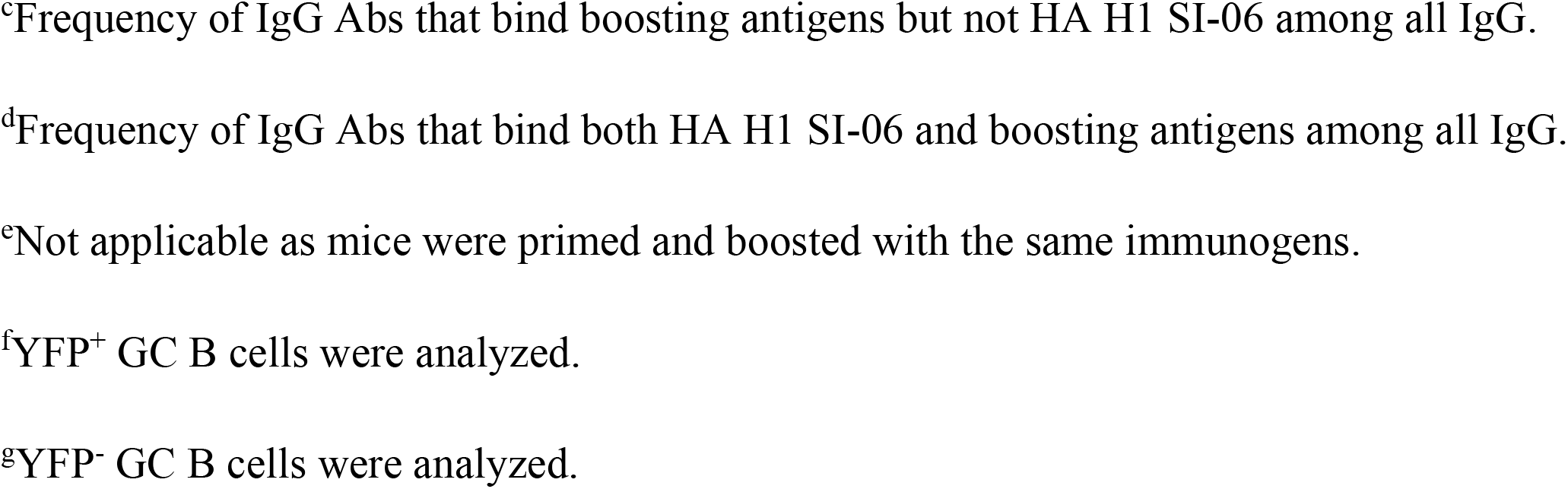
Summary of Single B Cell Cultures.

| Mouse/Treatment | Secondary Immunogens | Route of Boosts (Ipsi or Contra) | No. IgG Samples<sup>a</sup> | Primary Antigen-only<sup>b</sup> | Secondary Antigen-only<sup>c</sup> | Primary and Secondary Antigens<sup>d</sup> |
| --- | --- | --- | --- | --- | --- | --- |
| B6 | HA H1 SI-06 | Ipsi | 2,255 | NA<sup>e</sup> | NA<sup>e</sup> | 16% |
| B6 | HA H1 SI-06 | Contra | 2,858 | NA<sup>e</sup> | NA<sup>e</sup> | 7.5% |
| AID-Cre/EYFP Ctrl IgG or None | HA H1 SI-06 | Ipsi, YFP<sup>+</sup><sup>f</sup> | 514 | NA<sup>e</sup> | NA<sup>e</sup> | 21% |
| AID-Cre/EYFP Ctrl IgG or None | HA H1 SI-06 | Ipsi, YFP<sup>-g</sup> | 1,014 | NA<sup>e</sup> | NA<sup>e</sup> | 12% |
| AID-Cre/EYFP MR-1 Ab | HA H1 SI-06 | Ipsi, YFP<sup>+</sup><sup>f</sup> | 343 | NA<sup>e</sup> | NA<sup>e</sup> | 21% |
| B6 | HA H3 X31 | Ipsi | 1,825 | 1.2% | 12% | 1% |
| B6 | HA H3 X31 | Contra | 1,696 | 0.6% | 17% | 0.2% |
| B6 | rPA | Ipsi | 1,130 | 1% | 25% | 0.4% |
| B6 | rPA | Contra | 1,082 | 0.1% | 17% | 0% |

<sup>a</sup>Number of IgG<sup>+</sup> single B-cell cultures analyzed for each group.
<sup>b</sup>Frequency of IgG Abs that bind HA H1 SI-06 but not boosting antigens among all IgG.
<sup>c</sup>Frequency of IgG Abs that bind boosting antigens but not HA H1 SI-06 among all IgG.
<sup>d</sup>Frequency of IgG Abs that bind both HA H1 SI-06 and boosting antigens among all IgG.
<sup>e</sup>Not applicable as mice were primed and boosted with the same immunogens.
<sup>f</sup>YFP<sup>+</sup> GC B cells were analyzed.
<sup>g</sup>YFP<sup>-</sup> GC B cells were analyzed.

To compare the distribution of BCR affinities for HA H1 SI-06, we determined the avidity index (AvIn) for each clonal IgG Ab (*25*). The median AvIn for the GC B cells following ipsilateral boosts was 8 × 10^−4^, which was moderately but significantly different from median AvIn value for their contralateral boost counterparts (median AvIn = 5 × 10^−4^; *p* = 0.019; Fig. 2C). In addition, ipsilateral secondary GCs contained more high avidity (AvIn > 0.1) B cells. High avidity B cells constituted 14% (±14%) of HA-specific GC B cells elicited by ipsilateral boosts, about 4-fold above that of contralateral boost GC B cells (*p* = 0.035; Fig. 2D). Secondary GCs after local boosts were similar in size to those induced at distal sites but contained more high affinity B-cell clones.

### SHM in secondary GC B cells

To examine somatic genetics of secondary GC B cells, we determined VDJ gene sequences for subsets of Nojima culture samples (ipsilateral boosts, n = 837; contralateral boosts, n = 376). We amplified VDJ rearrangements from cDNA prepared and sequenced from the cell pellets of individual IgG^+^ cultures (*25*). Secondary GC B cells following ipsilateral boosts and contralateral boosts carried on average 2.7 and 2.0 V_H_ mutations, respectively (Fig. 3A), corresponding to V_H_ mutation frequencies (number of nucleotide substitutions per base pairs sequenced) of 1.0% and 0.7%, respectively (*p* = 0.73; Fig. 3A). These mutation frequencies were significantly (*p* < 0.001) higher than those in day 8 primary GC B cells (average 0.5%, Fig. 3A and (*25*)). Despite similar V_H_ mutation frequencies, secondary GC B cells following ipsilateral boosts had a broader range of V_H_ mutations (0-29) than their contralateral-boost counterparts [0-14 (Fig. 3A)]; 7.3% of secondary GC B cells following ipsilateral boosts carried 8 or more V_H_ mutations, while <1% of their contralateral-boost counterparts did (*p* < 0.001, Fig. 3B).

**Figure 3.**
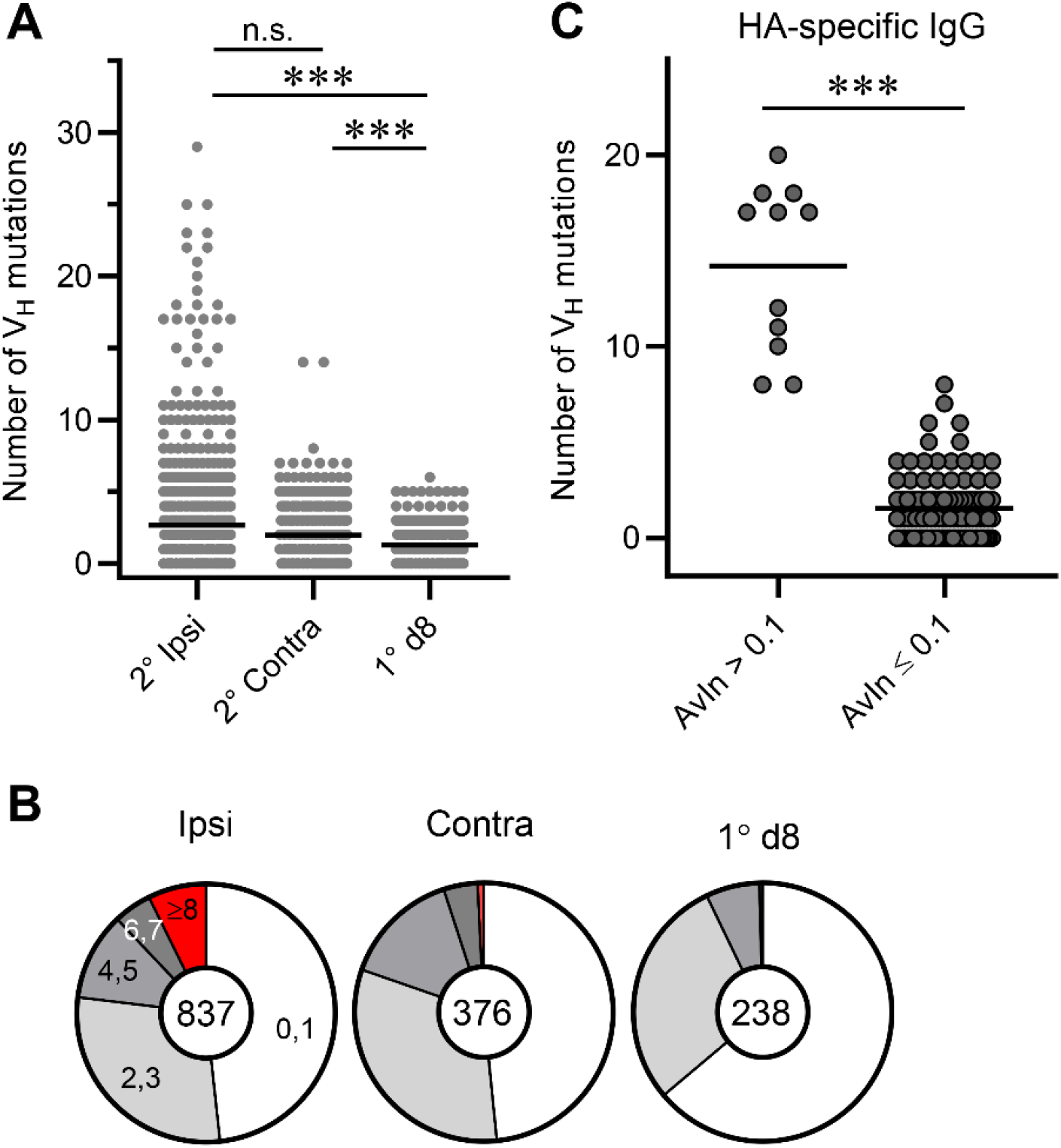
SHM in secondary GC B cells. Rearranged V_H_ gene segments in Nojima cultures were amplified from cell pellets of individual wells and V_H_ point mutations from germline enumerated. (**A**) Distribution of the number of V_H_ point mutations recovered from secondary GC B cells following ipsilateral boosts (n = 837) and distal boosts (n = 376). Data for primary d8 GC B cells (n = 238) are re-plotted from published work of our own (*25*). Each dot represents an individual IgG^+^ sample. Horizontal bars represent mean. ***, *p* < 0.001; n.s., *p* > 0.05 by Kruskal-Walis test with Dunn’s multiple comparisons. (**B**) Pie charts depict proportion of IgG examples that carry indicated number of V_H_ point mutations. (**C**) V_H_ gene sequences for HA H1 SI-06 reactive IgGs were split by their AvIn values into high avidity (AvIn > 0.1; n = 11) and the rest (AvIn ≤ 0.1; n = 102). ***, *p* < 0.001 by Mann-Whitney’s U test.

High avidity (AvIn > 0.1), HA-reactive IgG Abs recovered from secondary GC B cells carried elevated number of V_H_ mutations (range 8-20) (Fig. 3C), suggesting that these cells originated from the progeny of primary GC B cells rather than from newly-activated, naïve B cells (*11*). Elevated frequencies, in secondary GCs following ipsilateral boosts, of B cells that carry a large number of V_H_ mutations suggest an efficient engagement of the progeny of primary GC B cells at local sites.

### Progeny of the primary GC B cells enter secondary GCs more efficiently after local rather than distal boosts

Secondary GCs contain both the re-activated, progeny of primary GC B cells (*e*.*g*., persistent GC B and recalled Bmem cells) and newly-activated naïve B cells. Although the latter B-cell type dominates the response, at least in mice (*11*), elevated frequencies of higher affinity B cells and of highly-mutated B cells in secondary GCs following ipsilateral boosts (Figs. 2 and 3) suggest that B cells of the former types participate more efficiently in secondary GCs elicited by ipsilateral boosts. To directly compare ipsilateral and contralateral boosts for their efficiency in activating/recruiting the progeny of primary GC B cells into secondary GC responses, we used the AID-Cre-EYFP mouse model in which we labeled GC B cells during primary responses and traced the fates of the labeled B cells following boost immunizations (*1, 11*). We immunized AID-Cre-EYFP mice in the footpad with HA H1 SI-06, and then induced YFP expression in AID-expressing GC B cells by *i*.*p*. injections of tamoxifen (days 8-12). Eight-to ten weeks after primary immunizations, we boosted these animals with homologous HAs in the hock either ipsilaterally or contralaterally and enumerated frequency and number of YFP^+^ cells in PBs/PCs and GC B cell compartments in the draining LNs eight days following boosts.

Tamoxifen injections can induce YFP expression not only in AID-expressing cells that are activated by primary immunizations but also in those that are irrelevant to the priming (*e*.*g*., Peyer’s patch GC B cells constitutively present in the gut). To ensure that YFP^+^ cells that engage in secondary responses originated from cells specifically generated by the priming, we compared frequencies of YFP^+^ cells following “boost” immunizations with or without prior immunization. In the absence of priming, “boost” immunizations following tamoxifen injections recruited little or no YFP^+^ cells into PBs/PCs and GC B-cell compartments while YFP^+^ cells were readily detectable in these B-cell compartments in mice that had been primed previously (Fig. S3A). We conclude that most, if not all, YFP^+^ PBs/PCs and GC B cells present in draining LNs following boost immunizations are progeny of primary GC B cells elicited by the footpad immunization.

Consistent with comparable levels of recall serum IgG responses following ipsilateral and contralateral boosts (Fig. 1), these boost regimens elicited nearly identical frequencies and numbers of YFP^+^ PBs/PCs in the draining LNs (Fig. S3). Frequencies of YFP^+^ cells among PBs/PCs were 19% (±12%) and 18% (±9.6%) following ipsilateral and contralateral boosts, respectively (*p* > 0.99; Figs. S3B and S3C). Similarly, there were no significant differences in the number of YFP^+^ PBs/PCs in the draining LNs following ipsilateral and contralateral boosts (2,600 ±2,900 and 2,000 ±2,700, respectively; *p* > 0.99; Fig. S3D).

In contrast to plasmacytic responses, but consistent with overrepresentation of both high affinity B cells and highly-mutated B cells in secondary GCs (Figs. 2 and 3), the levels of YFP^+^ GC B cells in the draining LNs following ipsilateral boosts were higher than those following contralateral boosts. The frequency and number of YFP^+^ GC B cells after ipsilateral boosts were, respectively, ∼3-fold and ∼5-fold higher than those of their contralateral counterparts (frequency, 6.7% ±5.8% vs. 2.2% ±2%, *p* = 0.028; cell number, 7,000 ±4,400 vs. 1,200 ±1,100, *p* = 0.0038; Figs 4A-4C). We obtained similar results with *S1pr2*-ERT2cre-tdTomato mice, in which labeling efficiency of primary GC B cells is higher than that of AID-Cre-EYFP mice [85% vs. 25%; (*1, 17, 26*) and Fig. S4A)]. With *S1pr2*-ERT2cre-tdTomato mice, the frequency and number of labeled, secondary GC B cells was significantly higher in ipsilateral boosts [14% (±0.6%) vs. 4.8% (±4.3%); cell numbers, 22,000 (±9,500) vs. 5,000 (±1,600); Figs. S4B and S4C]. Local boosts activate and recruit the progeny of primary GC B cells into secondary GC responses more efficiently than do distal boosts.

**Figure 4.**
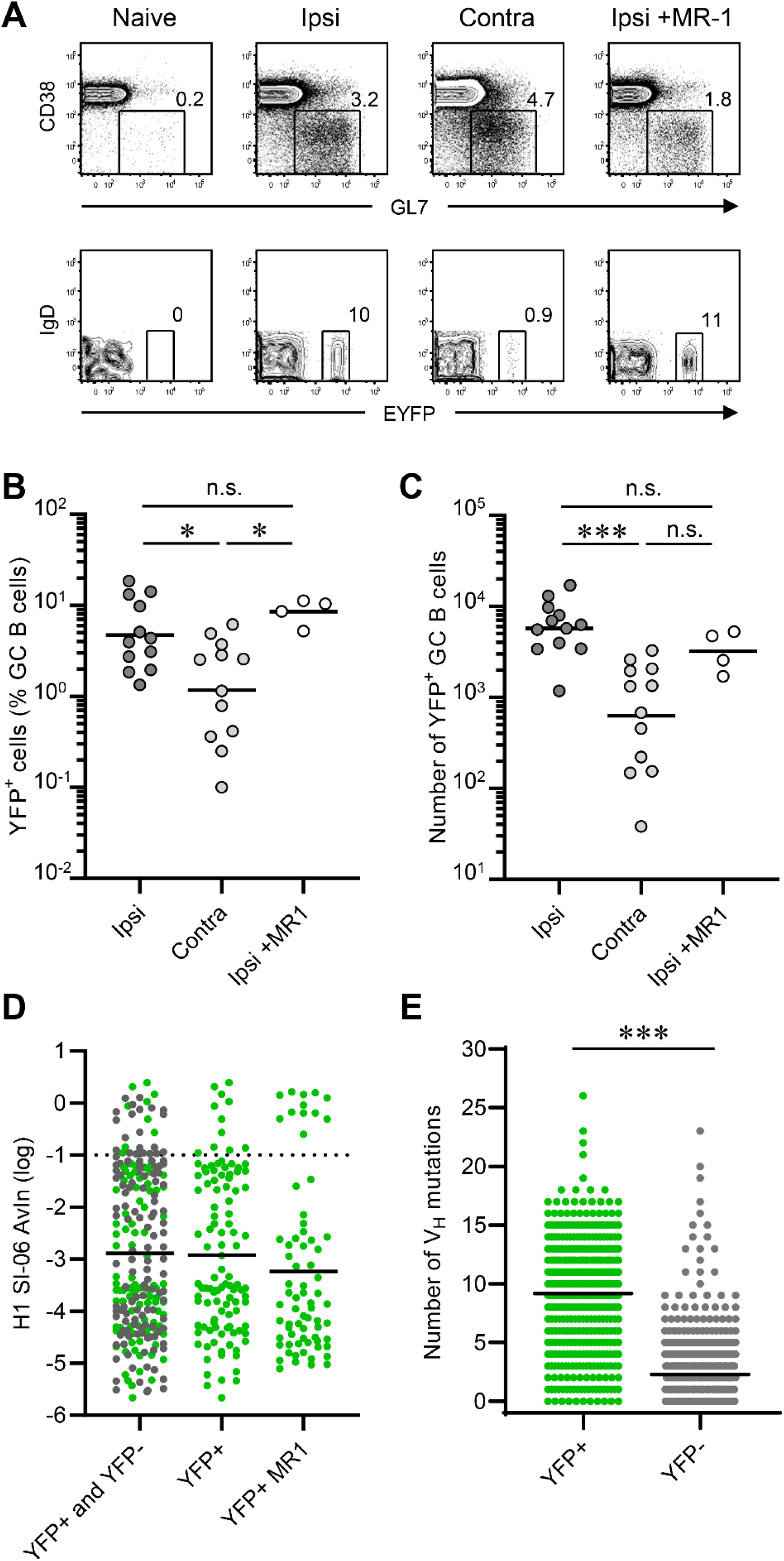
Progeny of the primary GC B cells engage in secondary GCs more efficiently after local boosts than after distal boosts. AID-Cre-EYFP mice were primed and boosted with H1 SI-06 as shown in Fig. 1A. Primed animals were given tamoxifen *i*.*p*. daily through days 8-12 after priming. In some experiments, primed animals received MR-1 Abs or control IgGs 4 weeks after the priming. Representative flow diagrams (**A**), and frequency (**B**) and number (**C**) of YFP^+^ GC B cells in the draining LNs are shown. (**B** and **C**) Each dot represents an individual mouse. Horizontal bars represent geometric mean. ***, *p* < 0.001; *, *p* < 0.05; n.s., *p* > 0.05 by Kruskal-Walis test with Dunn’s multiple comparisons. (**D**) Distributions of AvIn values for clonal IgGs from GC B cells following ipsilateral boosts are shown (left, YFP^+^ and YFP^-^ combined, n = 227; middle, YFP^+^, n = 109; right, YFP^+^ receiving MR-1, n = 71). Horizontal bars represent geometric mean. Dotted line represents AvIn = 0.1, our cutoff for “high avidity”. Combined data from 4 independent experiments are shown. No statistical significance was seen among groups by Kruskal-Walis test with Dunn’s multiple comparisons. (**E**) Distributions of the number of V_H_ point mutations in YFP^+^ (n = 399) and YFP^-^ (n = 618) secondary GC B cells following ipsilateral boosts. Each dot represents an individual IgG^+^ sample. Horizontal bars represent mean. ***, *p* < 0.001 by Mann-Whitney’s U test.

### Late disruption of primary GCs does not impair recruitment of their progeny to local boost GCs

Although GC responses are generally thought to be short-lived, persistent GCs following immunization have been reported (*1, 3, 27, 28*). Indeed, we observed small but detectable levels of GC B cells by flow cytometry for up to ∼2-3 months after primary immunizations (Figs. 2A, 2B and Figs. S2A-S2C). Therefore, in all preceding experiments, ipsilateral boosts may have (re)activated residual, primary GC B cells in addition to Bmem cells. To determine whether disruption of ongoing primary GC responses would reduce the engagement of YFP^+^ cells in secondary GCs, we injected a cohort of H1-primed AID-Cre-EYFP mice with CD154-specific MR-1 Ab or control hamster IgG 4 weeks after the priming (*29-31*), and then boosted these mice with H1 SI-06 4-6 weeks later (*i*.*e*., 8-10 weeks after priming). Eight days after boosts, we enumerated the frequency and number of YFP^+^ GC B cells.

Flow cytometric analysis performed at 5 weeks after MR-1 Ab injections (*i*.*e*., 9 weeks after priming, without boosts) showed that the number of YFP^+^ GC B cells was lower (∼30%) than in the control IgG group (580 vs. 1,900; Fig. S3E). Consistent with flow cytometric analysis, we observed in the draining LNs of control mice clusters of YFP^+^IgD^-^ cells in association with the CD21/CD35^bright^ follicular dendritic cell (FDC) network (Fig. S3F). In the MR-1 treated mouse LNs, FDC areas were filled with IgD^+^ B cells and only rarely were YFP^+^IgD^-^ cells detected (Fig. S3F), confirming that MR-1 injections had effectively disrupted the primary GC response. Despite absence of organized GC structures in MR-1-treated LNs, the majority (≥70%) of YFP^+^B220^+^CD138^-^ cells were GL-7^+^CD38^lo^, a surface phenotype indistinguishable from that of GC B cells (Fig. S3G).

Disruption of primary GC responses by CD154 blockade did not impair engagement of YFP^+^ cells in secondary GCs. Following ipsilateral boosts, the frequency and number of YFP^+^GC B cells were no different between mice that had received control IgGs and MR-1 Abs (Figs 4A-4C). These results suggest that progeny of the primary GC B cells that are resistant to the blockade of CD40:CD154 signaling represent the majority of YFP^+^ B cells in secondary GCs following ipsilateral boosts.

### Progeny of the primary GC B cells with a broad range of BCR affinities and of V_H_ SHM engage in secondary GCs

To examine the BCR repertoire of YFP^+^ secondary GC B cells, we sorted them following ipsilateral boosts, performed single-cell Nojima cultures and determined binding specificity and avidity of the clonal IgG Abs by a Luminex multiplex assay. Of the 514 clonal IgG Abs we recovered from the YFP^+^ secondary GC B cells, 109 (21%) bound to HA H1 SI-06 (Table 1). These HA-reactive clonal IgG Abs had a broad range of BCR affinities for the HA immunogen. This avidity distribution was no different from that of overall secondary GC B cells (compare Fig. 2C, Fig. 4D left column, and Fig. 4D middle column). Suppression of primary GC responses by the MR-1 treatment did not change the BCR avidity distribution of the YFP^+^ secondary GC B cells (Fig. 4D right column).

YFP^+^ secondary GC B cells, like all secondary GC B cells (Fig. 3A), had a broad range of V_H_ SHM (0-26; Fig. 4E). Consistent with a primary GC B-cell origin, YFP^+^ secondary GC B cells had higher V_H_ SHM (median number = 10; Fig. 4E) than did primary newly activated, d8 GC B cells (median = 1; Fig. 3A). In contrast to YFP^+^ cells, YFP^-^ secondary GC B cells included both recently-activated naïve B cells and progeny of primary GC B cells that were not labeled by tamoxifen ((*1, 26*) and Fig. S4). Unlike primary d8 GC B cells (Fig. 3A), highly-mutated clones (V_H_ mutations ≥8) constituted ∼5% (31/618) of the YFP^-^ secondary GC B cells (Fig. 4E). While these highly-mutated clones likely originated from the progeny of primary GC B cells, comparable levels of overall V_H_ SHM between the YFP^-^ secondary GC B cells and primary d8 GC B cells (median = 1, Figs. 3A and 4E) were in line with an estimation – most of the YFP^-^ secondary GC B cells originated from recently-activated naïve B cells. Both HA-binding and non-binding YFP^+^ secondary GC B cells had higher V_H_ SHM than did their respective YFP^-^ counterparts (Figs. S5A and S5B). Although high avidity (AvIn > 0.1), HA-binding YFP^+^ cells had elevated V_H_ SHM (range: 9-15), neither the number of V_H_ mutations nor of amino acid substitutions was correlated with AvIn values (Figs. S5C and S5D). Thus, progeny of primary GC B cells with a broad range of BCR affinities and of V_H_ SHM, including high affinity clones and highly-mutated clones, engage in secondary GC responses following ipsilateral boosts with homologous antigens.

### Recall of H1-specific serum IgG following local boosts with H3 HAs in H1-primed mice

To determine whether locality had any impact on recall humoral responses following boosts with heterologous antigens, we immunized B6 mice with HA H1 SI-06 in the right footpad, and then boosted these animals 8-12 weeks later in the hock ipsilaterally or contralaterally with a heterologous HA (HA H3 X31) or with an irrelevant antigen (recombinant protective antigen from *Bacillus Anthracis*, rPA). Eight days after boosts, we collected serum samples and draining LNs for quantifying HA-specific IgG Abs and for characterizing GC B cells, respectively (Fig. 5A).

**Figure 5.**
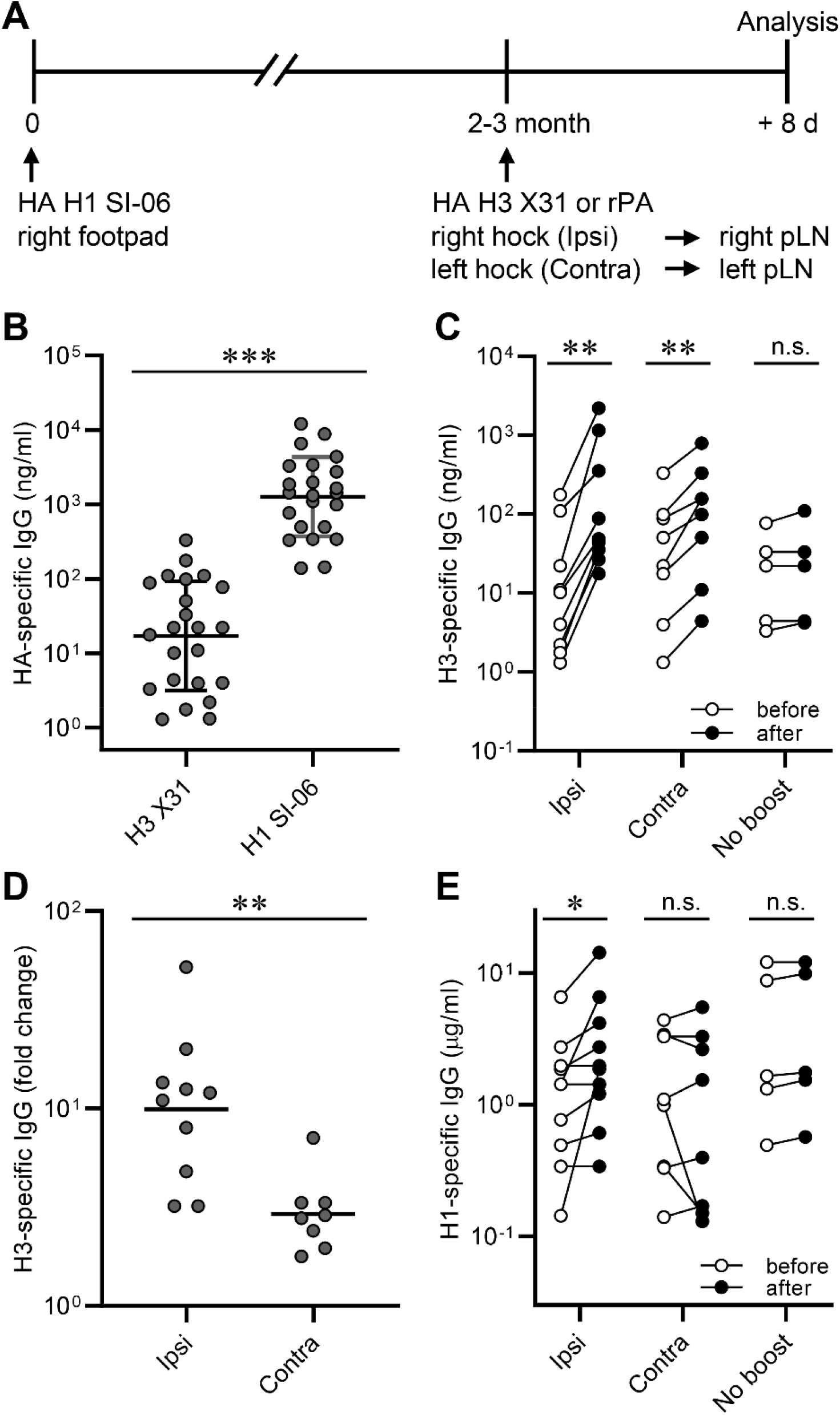
Recall of H1 HA-binding IgG Abs following local boosts with H3 HAs in H1-primed mice. Serum IgG responses following boosts with heterologous antigens. (**A**) Graphic representation of the immunization strategy used in experiments described in Figs. 5 and 6. (**B**) Concentrations of serum IgGs binding to HA H1 SI-06 and HA H3 X31 before boosts (n = 23). ***, *p* < 0.001 by Mann-Whitney’s U test. (**C**) Concentrations of serum IgGs specific to HA H3 X31 before and after boosts ipsilaterally (n = 10) or contralaterally (n = 8). No-boost group (n = 5) received primary immunizations but not boosts. Each symbol represents an individual mouse, and serum samples from the same animals are connected with lines. **, *p* < 0.01; n.s., *p* > 0.05 by Wilcoxon matched-pairs signed rank test. (**D**) Fold changes in concentrations of H3 HA-specific IgGs in serum samples in (**C**) after boosts. **, *p* < 0.01; by Mann-Whitney’s U test. (**E**) Concentrations of serum IgGs specific to HA H1 SI-06 in serum samples in (**C**). Each symbol represents an individual mouse, and serum samples from the same animals are connected with lines. *, *p* < 0.05; n.s., *p* > 0.05 by Wilcoxon matched-pairs signed rank test. Combined data from 4 independent experiments are shown.

Before boosting, sera from H1-primed animals contained H3-binding serum IgG Abs at 17 ng/ml (geometric mean), ∼70-fold lower than the concentration of H1-bindig IgG Abs (1,300 ng/ml; Fig. 5B). Following boosts, concentrations of H3-specific serum IgG Abs increased by 10-fold (from 13 ng/ml to 130 ng/ml, ipsilateral boosts) and 3-fold (from 27 ng/ml to 79 ng/ml, contralateral boosts; Fig. 5C and 5D). In contrast to H3-binding IgG responses, ipsilateral, but not contralateral, boosts with H3 HAs increased concentrations of H1-specific serum IgG Abs by 2-fold (from 1.1 μg/ml to 2.1 μg/ml, *p* < 0.05; Fig. 5D), implying that ipsilateral boosts with HA H3 X31 re-activated H1/H3 cross-reactive cells that had been established by priming with HA H1 SI-06 more efficiently than did contralateral boosts.

### More H1/H3 cross-reactive secondary GC B cells after local boosts with H3 HA in H1-primed mice

We determined the number and frequency of secondary GC B cells by flow cytometry. Like secondary GC responses to homologous HA boosts (Fig. 2A), boosts with heterologous HAs elicited robust GC responses in the draining LNs following both ipsilateral and contralateral boosts (Fig. 6A, 6B). To determine the reactivity profile of secondary GC B cells after boosting with heterologous HA antigens, we isolated GC B cells from the draining LNs of boosted animals and established single-cell Nojima cultures (*25*). On average, antigen-specific B cells (*i*.*e*., clonal IgGs that bound HAs of H1 SI-06 or H3 X31 or both) accounted for 16% (±9.4%) and 18% (±8.2%) of clonal IgGs recovered from secondary GC B cells following ipsilateral and contralateral boosts, respectively (Fig. S6A). For both boost regimens, most (90% and 97%, ipsilateral and contralateral boosts, respectively) HA reactive B cells bound HA H3 X31 (*i*.*e*., the boosting antigen; Fig. 6C). Despite these similarities, the distribution of HA reactivity of secondary GC B cells was different between boost regimens. The frequency of H1/H3 cross-reactive cells among HA-specific secondary GC B cells was significantly higher after ipsilateral boosts than after contralateral boosts (Fig. 6C, 6D). On average, H1/H3 cross-reactive B cells constituted 12% (±11%) and 0.9% (±1.5%) of HA-reactive GC B cells after ipsilateral and contralateral boosts, respectively. We recovered H1/H3 cross-reactive B cells from 8 out of 10 mice given an ipsilateral boost and 2 out of 6 mice after a contralateral boost (Fig. 6D).

The participation of cross-reactive GC B cells that bound both priming antigens and boosting antigens was much lower when we boosted H1 SI-06 primed mice with an irrelevant antigen, rPA. Antigen-specific B cells (*i*.*e*., clonal IgGs that bound H1 SI-06 or rPA or both antigens) accounted for 28% (±10%) and 18% (±3.3%) of clonal IgGs recovered from secondary GC B cells following ipsilateral boosts and contralateral boosts, respectively (Fig. S6B).

**Figure 6.**
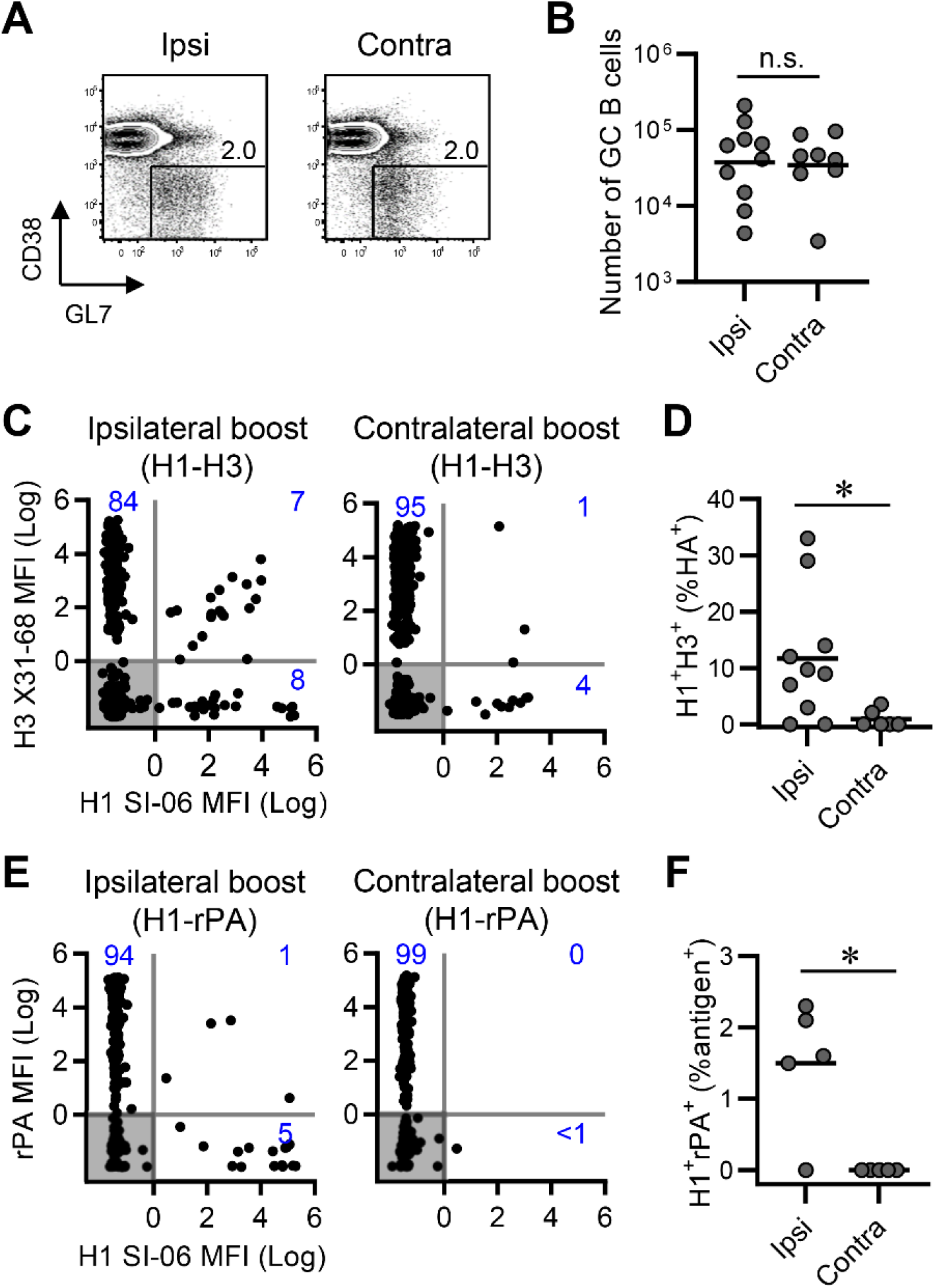
Participation of H1/H3 cross-reactive B cells in secondary GCs following local boosts with H3 HAs in H1 HA-primed mice. GC responses following boosts with heterologous antigens (See also a legend of Fig. 5A). (**A** and **B**) Representative flow diagrams of GL-7 and CD38 expressions on B220^+^CD138^-^ cells (**A**) and number of B220^+^CD138^-^GL-7^+^CD38^lo^IgD^-^ GC B cells in the draining LNs of boosted mice (ipsilateral boosts, n = 10; contralateral boosts, n = 8). Numbers near boxes in (**A**) represent frequency of GL-7^+^CD38^lo^IgD^-^ cells among B220^hi^CD138^-/lo^ cells. Horizontal bars in (**B**) represent geometric mean. n.s., *p* > 0.05 by Mann-Whitney’s U test. Combined data from 4 independent experiments are shown. (**C**-**F**) Individual GC B cells were placed in Nojima cultures for determination of BCR reactivity. H3 X31 ipsilateral boosts (n = 1,825), H3 X31 contralateral boosts (n = 1,696), rPA ipsilateral boosts (n = 1,130), and rPA contralateral boosts (n = 1,082). (**C** and **E**) Background-subtracted, normalized MFI values (Log_10_) for H1 SI-06 (x-axes) and H3 X31 (**C**, y-axes) or rPA (**E**, y-axes). Numbers in each quadrant represent frequencies of H1 or H3 or H1/H3 (**C**) reactive IgGs and H1 or rPA or H1/rPA reactive IgGs among antigen-specific clonal IgGs. Each dot represents an individual clonal IgG. (**D** and **F**) Frequencies of H1/H3 cross-reactive IgGs (**D**) and of H1/rPA cross-reactive IgGs (**F**) among all antigen-specific IgG. Each dot represents an individual mouse. *, *p* < 0.05 by Mann-Whitney’s U test. Combined data from 2 (rPA boosts) and 4 (H3 X31 boosts) independent experiments are shown.

Following ipsilateral boosts with rPA, H1/rPA yielded cross-reactive cells that represented 1% of antigen-specific B cells (Figs. 6E and 6F), and contralateral boosts with rPA yielded none (Figs. 6E and 6F). Epitopic similarity between the priming antigens and boosting antigens is thus a key determinant for cross-reactive B cells to engage in secondary GC responses.

## DISCUSSION

We have shown an important role of locality in secondary GC responses. Using a combined approach of single B-cell cultures with a fate-mapping mouse model, we found that in response to homologous HA antigens, ipsilateral boosts recruited to secondary GCs progeny of the primary GC B cells more efficiently than did contralateral boosts (Fig. 4). Consequently, secondary GCs following ipsilateral boosts comprised B cells with elevated IgH SHM and with high avidity to immunogens at higher frequencies than did their contralateral counterparts (Figs. 2-4). In response to heterologous HA antigens, ipsilateral boosts increased serum IgG that bound the priming HAs along with those that bound boosting HAs, while contralateral boosts increased only serum IgG components that bound boosting HAs (Fig. 5). Ipsilateral boosts recruited to secondary GCs cross-reactive B cells that bound both the priming and boosting HAs more efficiently than did contralateral boosts (Fig. 6).

Which factors contribute to an efficient engagement of the progeny of primary GC B cells in secondary GCs following ipsilateral boosts? Local antigens that are retained on FDCs (*32*) have significant roles in retaining “primed B cells” within the draining LNs without further differentiation into PBs/PCs and thereby contribute to long-lasting, local B-cell memory (*22-24*). We propose that local B-cell memory is retained in the primed LNs in the form of classical Bmem cells, persistent GC B cells, and GC-phenotype B cells that are independent of GC structures (Fig. S3) and that these persistent “primed B cells” contribute to recall GC responses at local sites. For example, “re-fueling” of persistent GCs with cognate antigens (*1, 3, 27, 28*)(Figs. 2, S2, and S3) may expand the persistent GC B cells following ipsilateral boosts (*11*). In addition to this re-fueling model, our observations show significant contributions of the MR-1 resistant B cells in secondary GC responses following local boosts, as the disruption of ongoing GC structures by the MR-1 Ab treatment did not impair participation of the fate-mapped, high affinity B cells in secondary GCs (Figs. 4 and S3). The majority (≥70%) of the MR-1 resistant, fate-mapped cells found in the primed LNs had surface molecule expressions that were indistinguishable from that of GC B cells (*i*.*e*., GL-7^+^CD38^lo^) despite the absence of organized GC structures (Fig. S3).

Our observations that ipsilateral boosts recruit to secondary GCs progeny of the primary GC B cells more efficiently than do contralateral boosts might provide a possible explanation for an apparent discrepancy between recent observations concerning Bmem recruitment to recall GCs. Mesin *et al*. (*11*) have suggested that recall of Bmem cells may be an inefficient process based on observations in fate-mapping mice. They find that fate-mapped cells (*i*.*e*., progeny of the primary GC B cells) are less frequent in secondary GCs following *contralateral* boosts. In contrast, Turner *et al*. (*4*) have analyzed the fine needle aspirates of the LN samples from recent influenza vaccinees and shown that a substantial fraction of antigen-binding GC B cells have elevated Ig SHM and that they are broadly cross-reactive, consistent with Bmem origin.

Although speculative, it is possible that the human donors studied by Turner *et al*. had established local B-cell memory from previous vaccinations and that *ipsilateral* boosts with an influenza vaccine activate local B-cell memory populations (described above), resulting in substantial engagement of Bmem cells in recall GCs.

Recall GCs following local boosts contain B cells of heterogeneous origin, including recently-activated, naïve B cells and recalled “primed B cells” – a category that comprises classical Bmem cells as well as persistent GC B cells and MR-1 resistant, GC-phenotype B cells (Fig. S3). What would be an outcome of the recall GC responses? That is, who will win the Darwinian selection that would be operated in recall GCs? While activated, naïve B cells that are overrepresented in early recall GCs (*11, 12*)(Fig. 4) may eventually outnumber the rarer, recalled “primed B cells” over the course of the GC reactions, a relative population size in early GCs does not necessarily predict an outcome of the B-cell selection. For example, a subset of recalled Bmem cells may have a “head start” and outcompete the naïve counterparts as they express high avidity BCRs that are infrequent in the naïve counterparts (Figs. 2-4). It is also possible that secondary GCs, like primary GCs, are permissive environments and overall BCR affinity maturation could operate without losing BCR diversity (*25*). In this model, both the rarer recalled “primed B cells” and more frequent, activated naïve B cells could co-exist throughout the recall GC responses and produce their respective progenies.

HA-binding IgGs accounted for only ∼20% of all clonal IgGs recovered from secondary GC B cells (Table 1). This frequency did not change when we focused our analysis on fate-mapped cells (Table 1). In other words, most (80%) of the progeny of primary GC B cells participate in secondary GC responses without measurable affinity to immunogens. One possible explanation would be that these HA non-binding B cells had lost affinity to HAs by new rounds of SHM during secondary GC responses. Indeed, without strong selection, any stochastic mutational process has a much greater likelihood of lowering BCR affinity than of increasing it. It is also possible that these HA non-binding YFP^+^ cells represent progeny of the antigen non-binding B cells found among primary GC B-cell populations (*25, 33*) and/or among the contemporaneously generated Bmem pool (*12*). Recovery of HA non-binding, YFP^+^ secondary GC B cells that carried no IgH SHM or only silent (no amino acid change) IgH SHM (Fig. S5) strongly suggests that they were activated by both the primary and secondary immunizations without measurable BCR affinities (as IgG) and yet participated in secondary GC responses.

Although the nature of these antigen non-binding secondary GC B cells still needs to be elucidated, secondary GCs, like primary GCs, are permissive environments, at least for the entry and early phase of the reaction, and support B cell clones with a broad range of BCR affinities, including those with undetectable affinities for the native form of the immunogen.

H1/H3 cross-reactive B cells participated in secondary GCs following ipsilateral boosts with H3 HAs, and to a lesser extent, following contralateral boosts in H1-primed mice (Fig. 6). This advanced engagement of H1/H3 cross-reactive B cells in local, secondary GCs accompanied an increase, following ipsilateral boosts, in binding of serum IgG Abs to H1 HAs as well as of those binding to H3 HAs (Fig. 5). Our observations suggest that local boosts with H3 HAs effectively activate H1/H3 cross-reactive B cells that are retained within the persistent pool of “primed B cells” we described above. Participation of H1/H3 cross-reactive B cells in secondary GCs would give them an opportunity to update their BCRs to fit with newly-introduced antigens (*i*.*e*., H3 HAs). A further question is how these cross-reactive B cells compete with other B cells (*e*.*g*., H1-specific or H3-specific) and how they evolve in recall GCs.

B-cell-lineage immunogen design vaccine strategies involve sequential immunizations with a series of designed immunogens to activate the targeted, naïve precursor B cells and their descendants to direct B-cell lineage progression toward a desired goal that would not be reached otherwise (*e*.*g*., broadly neutralizing Abs to HIV-1 or influenza) (*34*). As AID expression and SHM are largely limited in GC B cells (*35, 36*), one of the key steps is engagement of Bmem cells in recall GC responses in which they undergo new rounds of SHM, clonal selection and affinity maturation to newly-introduced antigens. Our observations suggest that activation of local B-cell memory populations by repeated immunizations at the same sites might be required to ensure efficient participation of “primed B cells” in recall GC responses, and thus might be necessary for a successful, B-cell-lineage immunogen design vaccine strategy.

## MATERIALS AND METHODS

### Mice and Immunizations

Female, C57BL/6 mice were obtained from the Jackson Laboratory. *Aicda*^Cre-ERT2^ x *Rosa26*^loxP-EYFP^ (AID-Cre-EYFP) mice (*1, 26*) and *S1pr2*-ERT2cre-tdTomato mice (*17*) were provided by Drs. Claude-Agnes Reynaud and Jean-Claud Weill and by Dr. Tomohiro Kurosaki, respectively. All mice were maintained under specific pathogen-free conditions at the Duke University

Animal Care Facility. Eight to 12-week-old mice were immunized with 20 μg of HA H1 SI-06 (see below) in the footpad of the right hind leg. One to three months later, cohorts of mice were boosted with 20 μg of HA SI-06 or HA H3 X31 or rPA (protective antigen from *Bacillus Anthracis*, BEI Resources) in the hock ipsilaterally (right hind leg) or contralaterally (left hind leg). Mice were analyzed 8 days after the boosts. All antigens were mixed with Alhydrogel^®^ before immunizations. In some experiments, we injected total 900 μg of MR-1 Abs or control hamster IgGs (300 μg daily for three consecutive days) *i*.*v*. 4 weeks after primary immunizations to disrupt ongoing GC reactions. LNs draining the site of the most recent immunizations (right popliteal LNs for no boosts and ipsilateral boosts, and left popliteal LNs for contralateral boosts) were analyzed. Sera were collected before boosts (one day before or same day as boosts) and after boosts (8 days after boosts). All experiments involving animals were approved by the Duke University Institutional Animal Care and Use Committee.

### Expression and purification of HAs

HAs for the full-length, soluble ectodomains of H1 A/Solomon Islands/03/2006 (H1 SI-06) and H3 A/Aichi/2/1968 (H3 X31) were cloned, expressed, and purified as previously described (*25, 37-39*).

### Flow Cytometry

GC B cells (GL-7^+^B220^hi^CD38^lo^IgD^-^CD138^-^) and plasmablasts/-cytes (B220^lo^CD138^hi^) in the draining LNs were identified as described (*25, 40, 41*). Labeled cells were analyzed/sorted in a FACS Canto (BD Bioscience) or a FACS LSRII (BD Biosciences) or a FACS Vantage with DIVA option (BD Bioscience). Flow cytometric data were analyzed with FlowJo software (Treestar Inc.). Doublets were excluded by FSC-A/FSC-H gating strategy. Cells that take up propidium iodide were excluded from our analyses.

### Single-cell Nojima culture

GC B cells were expanded in single cell cultures (*25*). Briefly, NB-21.2D9 cells feeder cells were seeded into 96-well plates at 2,000 cells/well in B cell media (BCM); RPMI-1640 (Invitrogen) supplemented with 10% HyClone FBS (Thermo scientific), 5.5 × 10^−5^ M 2-mercaptoethanol, 10 mM HEPES, 1 mM sodium pyruvate, 100 units/ml penicillin, 100 μg/ml streptomycin, and MEM nonessential amino acid (all Invitrogen). Next day, recombinant mouse IL-4 (Peprotech; 2 ng/ml) was added to the cultures, and then single B cells were directly sorted into each well of 96-well plates using a FACS Vantage. Two days after culture, 50% (vol.) of culture media were removed from cultures and 100% (vol.) of fresh BCM were added to the cultures. Two-thirds of the culture media were replaced with fresh BCM every day from day 4 to day 8. On day 10, culture supernatants were harvested for ELISA determinations and culture plates were stored at - 80°C for V(D)J amplifications.

### ELISA and Luminex assays

Presence of total and antigen-specific IgG in serum samples and in culture supernatants were determined by ELISA and Luminex multiplex assay (*25*). Diluted culture supernatants (1:10 in PBS containing 0.5% BSA and 0.1% Tween-20) were screened for the presence of IgGs by standard ELISA (*25*). IgG^+^ samples were then screened for binding to immunogen antigens (HA H1 SI-06, HA H3 X31, and rPA) by Luminex assay (*25*). Briefly, serum samples or culture supernatants were diluted (initial dilutions of 1: 100, and then 3-fold, 11 serial dilutions for serum samples; 1: 10 or 1: 100 for culture supernatants) in 1×PBS containing 1% BSA, 0.05% NaN3 and 0.05% Tween20 (assay buffer) with 1% milk and incubated for 2 hours with the mixture of antigen-coupled microsphere beads in 96-well filter bottom plates (Millipore). After washing with assay buffer, these beads were incubated for 1 hour with PE-conjugated goat anti-mouse IgG Abs (Southern Biotech). After three washes, the beads were re-suspended in assay buffer and the plates were read on a Bio-Plex 3D Suspension Array System (Bio-Rad). The following antigens were coupled with carboxylated beads (Luminex Corp): BSA (Affymetrix), goat anti-mouse Igκ, goat anti-mouse Igλ, goat anti-mouse IgG (all Southern Biotech), HA H1 SI-06, HA H3 X31, and rPA. For each IgG^+^ culture supernatant sample and serum sample, concentrations of antigen-binding IgG were determined in reference to monoclonal standards; “musinized” CH67 (*25*) and HC19 (*42, 43*) for H1-specific IgG and H3-specific IgG, respectively. Measured K_D_ of CH67 and HC19 IgG are 2.4-24 nM and 28 ±8 nM (*43*), respectively. AvIn values were obtained for HA H1 SI-06 binding IgG samples in reference to monoclonal standard, “musinized” CH67 (*25*).

### Amplification of V(D)J rearrangements

V(D)J rearrangements of cultured B cells were amplified by a semi-nested PCR. Total RNA was extracted from selected samples using Quick-RNA 96 kit (Zymo Research). cDNA was synthesized from DNase I-treated RNA using SMARTScribe™ Reverse Transcriptase (Clontech) with 0.2 μM each of gene-specific reverse primers (mIgGHGC-RT, mIgKC-RT, mIgLC23-RT, mIgLC1-RT, and mIgLC4-RT, Table S1) and 1 μM of 5’ SMART template-switching oligo that contained plate-associated barcodes (Table S1) at 50°C for 50 min followed by 85°C for 5 min. cDNA was then subjected to two rounds of PCR using Herculase II fusion DNA polymerase with combinations of forward primers and reverse primers that contained well-associated barcodes (Table S1). Primary PCR: 95°C for 4 min, followed by 2 cycles of 95°C for 30 sec, 64°C for 20 sec, 72°C for 30 sec; 3 cycles of 95°C for 30 sec, 62°C for 20 sec, 72°C for 30 sec; 25 cycles of 95°C for 30 sec, 55°C for 20 sec, 72°C for 30 sec; and 72°C for 10 min.

Secondary PCR: 95°C for 4 min, followed by 40 cycles of 95°C for 30 sec, 45°C for 20 sec, 72°C for 30 sec; and 72°C for 10 min. Gel-purified, pooled V(D)J amplicands were submitted to DNA Link, Inc. to obtain DNA sequences using PacBio sequencing platform. Obtained circular consensus sequences (CCS) were analyzed to build a consensus sequence for each sample. Obtained CCS were first sorted according to the barcode. Consensus sequences for individual samples were then built from 20 randomly-selected CCS (or from a minimum of 5 CCS) using established alignment programs MUSCLE v3.8.31 (*44*) and Kalign (*45*) with default settings. The rearranged V, D, and J gene segments were first identified using IMGT/V-QUEST (http://www.imgt.org/) or Cloanalyst (*46*), and then numbers and kinds of point mutations were determined.

### Immunofluorescence

Popliteal LNs from immunized AID-Cre/EYFP mice were fixed with 1% paraformaldehyde in PBS for overnight at 4°C, and then placed in PBS containing 10-30% sucrose at 4 °C (10% for 2 hours, 20% for 2 hours, and then 30% for overnight). Tissues were then embedded in Tissue-Tek O.C.T. compound (Sakura Finetek) and snap-frozen in 2-methylbutane cooled with liquid nitrogen. Frozen tissues were stored at -80 °C until use. Serial 5-μm-thick cryosections were cut on a CM1850 cryostat (Leica) and thaw-mounted onto glass slides and stored at -. After air drying, sections were rehydrated in a washing buffer (PBS with 0.5% BSA and 0.1% Tween-20) for 30 minutes at room temperature, and then blocked with rat anti-mouse CD16/32 (2.4G2) and rat IgG (Sigma-Aldrich) for 15 minutes at room temperature. Sections were labeled with AlexaFluor488-conjugated anti-GFP Ab (FM264G, BioLegend), BV510-conjugated anti-IgD (11-26c.2a, BioLegend), and AlexaFluor647-conjugated anti-CD21/35 (7E9, BioLegend) in a humidified chamber for 3 hours at room temperature in the dark. After washing, labeled sections were mounted in Fluoremount-G (SouthernBiotech) and imaged with SP8 upright confocal microscope (Leica). The images were processed with ImageJ software (Fiji package, NIH).

### Statistics

Statistical significance (*p* < 0.05) was determined by Wilcoxon matched-pairs signed rank test, Kruskal-Walis test with Dunn’s multiple comparisons, and Mann-Whitney’s *U* test using GraphPad Prism software (version 9.2.0, GraphPad Software). Statistic test is indicated within each figure legend.

## Supplementary Materials

**Figure S1.**
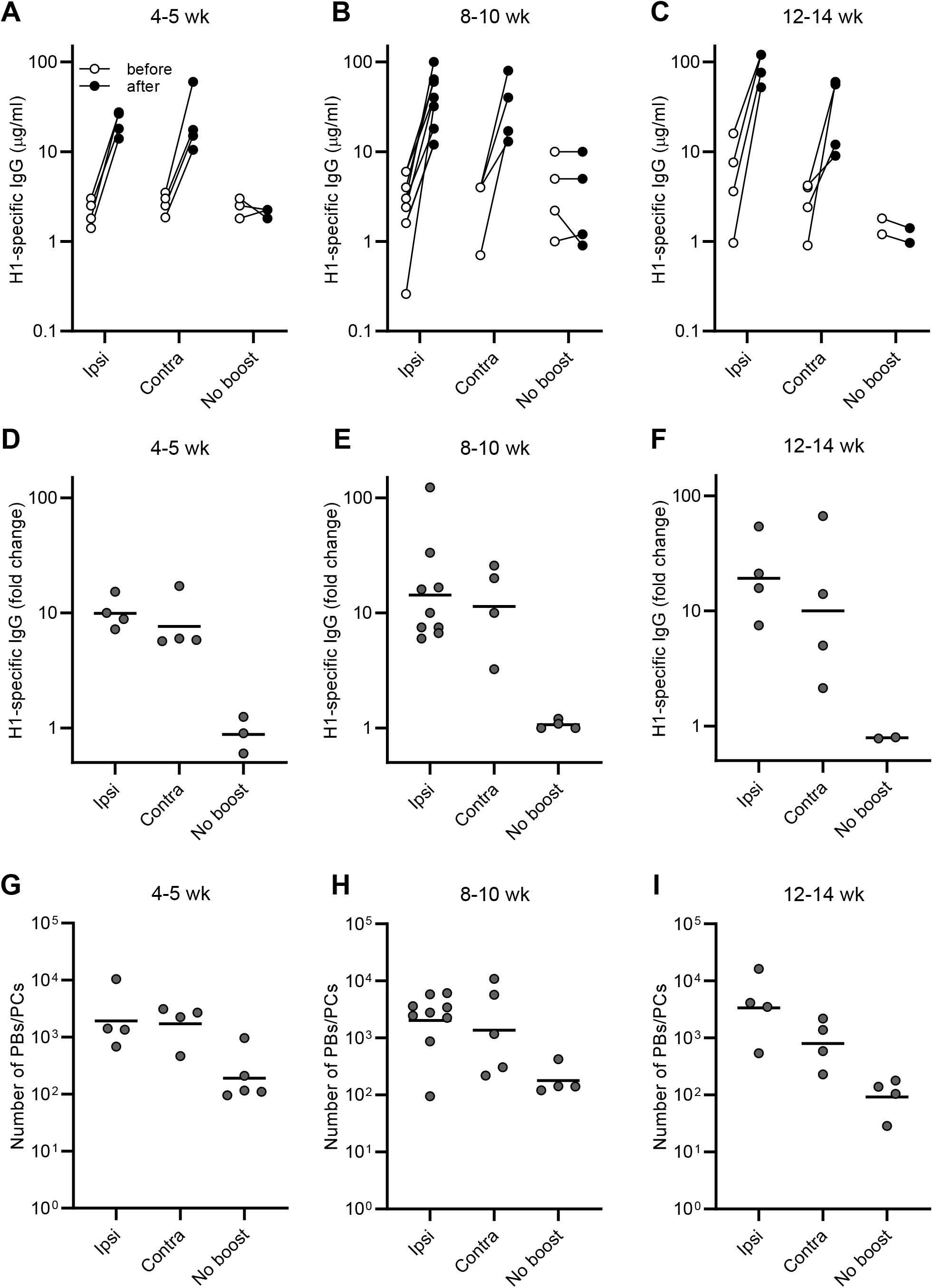
Robust recall antibody responses following boost immunizations. Data shown in Fig. 1 are split by the intervals between the priming and boosting (4-5 wk, 8-10 wk, and 12-14 wk). Concentrations of serum IgGs specific to H1 SI-06 (**A**-**C**), fold changes in concentrations of H1-specific IgGs in serum samples after boosts (**D**-**F**), and number of B220loCD138hi plasmablasts/cytes in the draining LNs (**G**-**I**) for indicated prime/boost intervals are shown. Each dot represents an individual serum sample or mouse. Horizontal bars in (**D**-**I**), geometric mean. See also figure legend for Fig. 1.

**Figure S2.**
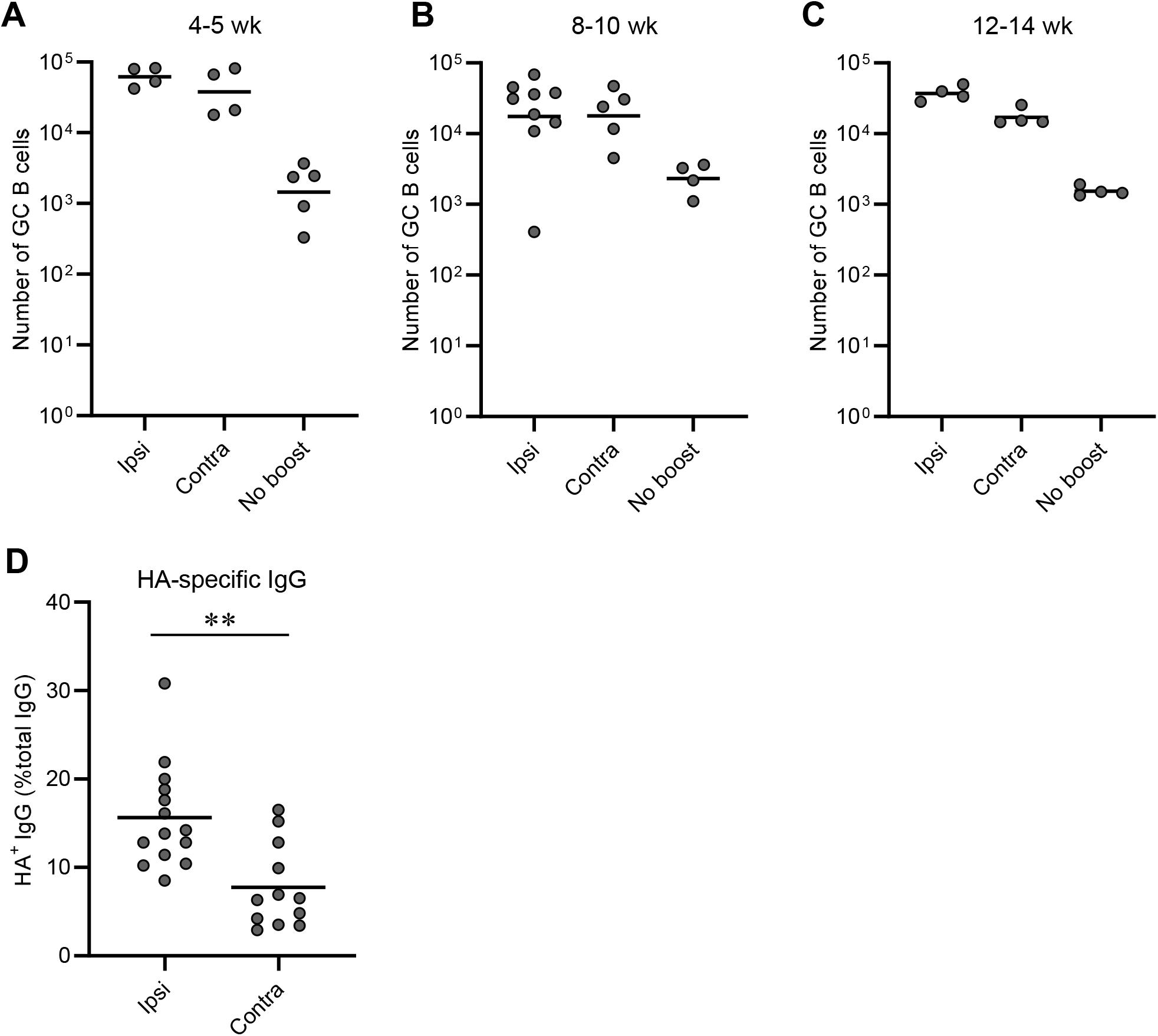
Robust GC responses following boosts at local sites or distal sites. (**A**-**C**) Data shown in Fig. 2B are split by the intervals between the priming and boosting (4-5 wk, 8-10 wk, and 12-14 wk), and number of B220+CD138-GL-7+CD38loIgD-GC B cells in the draining LNs for the indicated prime/boost intervals is shown. Each dot represents an individual mouse. Horizontal bars, geometric mean. See also figure legend for Fig. 2. (**D**) Frequency of HA-specific IgGs among all clonal IgGs for secondary GC B cells elicited by ipsilateral or contralateral boosts. Each dot represents an individual mouse. Horizontal bars, mean. *?*, *p* < 0.01 by Mann-Whitney’s U test.

**Figure S3.**
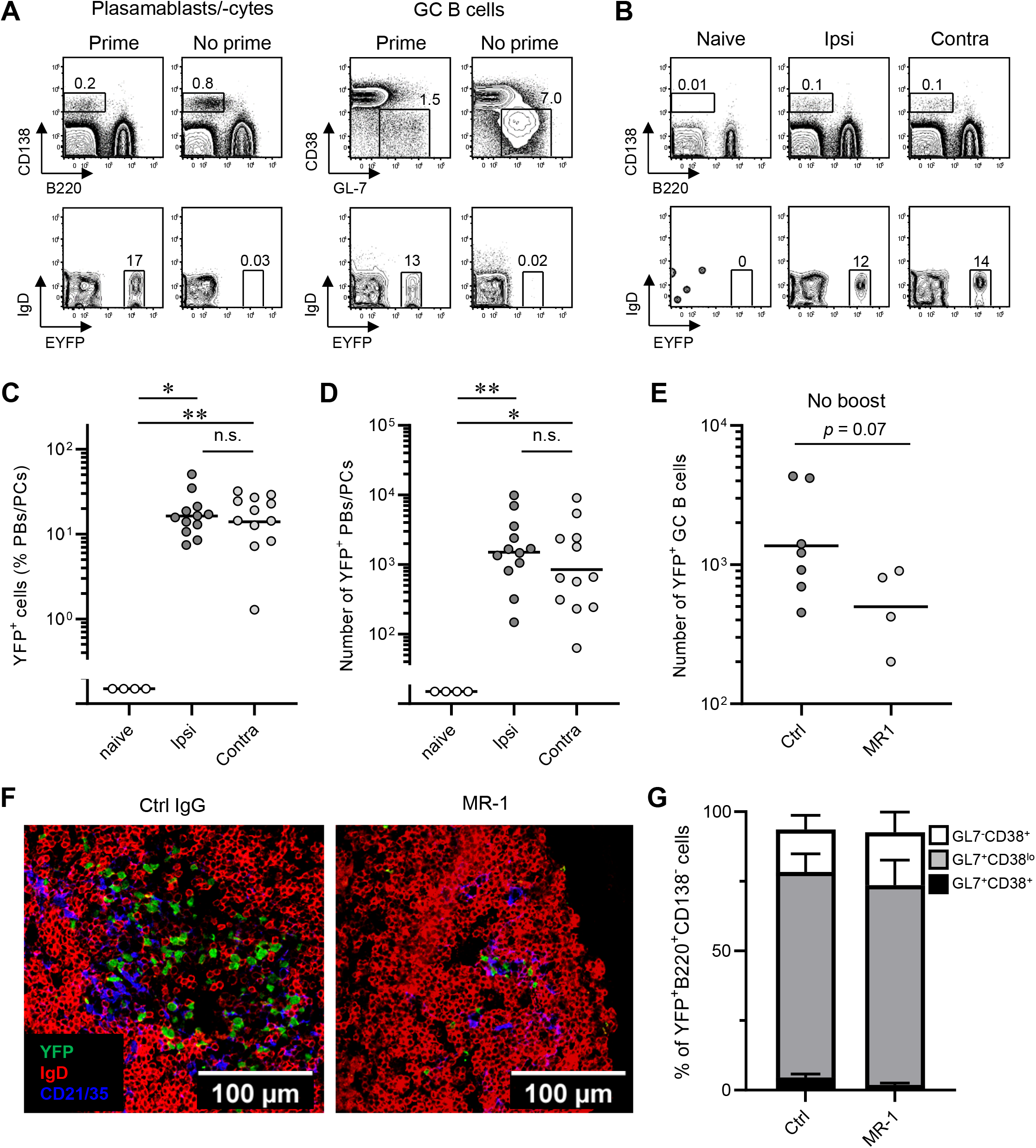
Tracing the fates of the progeny of primary GC B cells in AID-Cre-EYFP mice. AID-Cre-EYFP mice that had or had not received primary immunizations (Prime and No prime, respectively) received tamoxifen. These mice were then “boosted” ipsilaterally. Following boosts (d8), frequencies of YFP+ cells among PBs/PCs and GC B cells were determined by flow cytometry. (**B**-**D**) AID-Cre-EYFP mice were primed and boosted with H1 SI-06 (see also figure legend for Fig. 2). Representative flow diagrams (**B**), and frequency (**C**) and number (**D**) of YFP+ PBs/PCs in the draining LNs are shown. (**C** and **D**) Each dot represents an individual mouse. Horizontal bars, geometric mean. *, *p* < 0.05; **, *p* < 0.01; n.s., *p* > 0.05 by Kruskal-Walis test with Dunn’s multiple comparisons. (**E**-**G**) AID-Cre-EYFP mice that had received primary immunizations and tamoxifen were treated *i*.*v*. with control IgGs or MR-1 antibodies 4 weeks after the priming. Four to five weeks later, YFP+ B cells were assessed by flow cytometry (**E**) and immunofluorescence (**F**). Each dot in (**E**) represents an individual mouse. Horizontal bars, geometric mean. *p* = 0.07 by Mann-Whitney’s U test. (**G**) Proportion of GL-7-CD38+, GL-7+CD38-, and GL-7+CD38+ cells among all YFP+B220+CD138-cells. Error bars, SEM, n = 7 and 4 for Ctrl and MR-1 groups, respectively.

**Figure S4.**
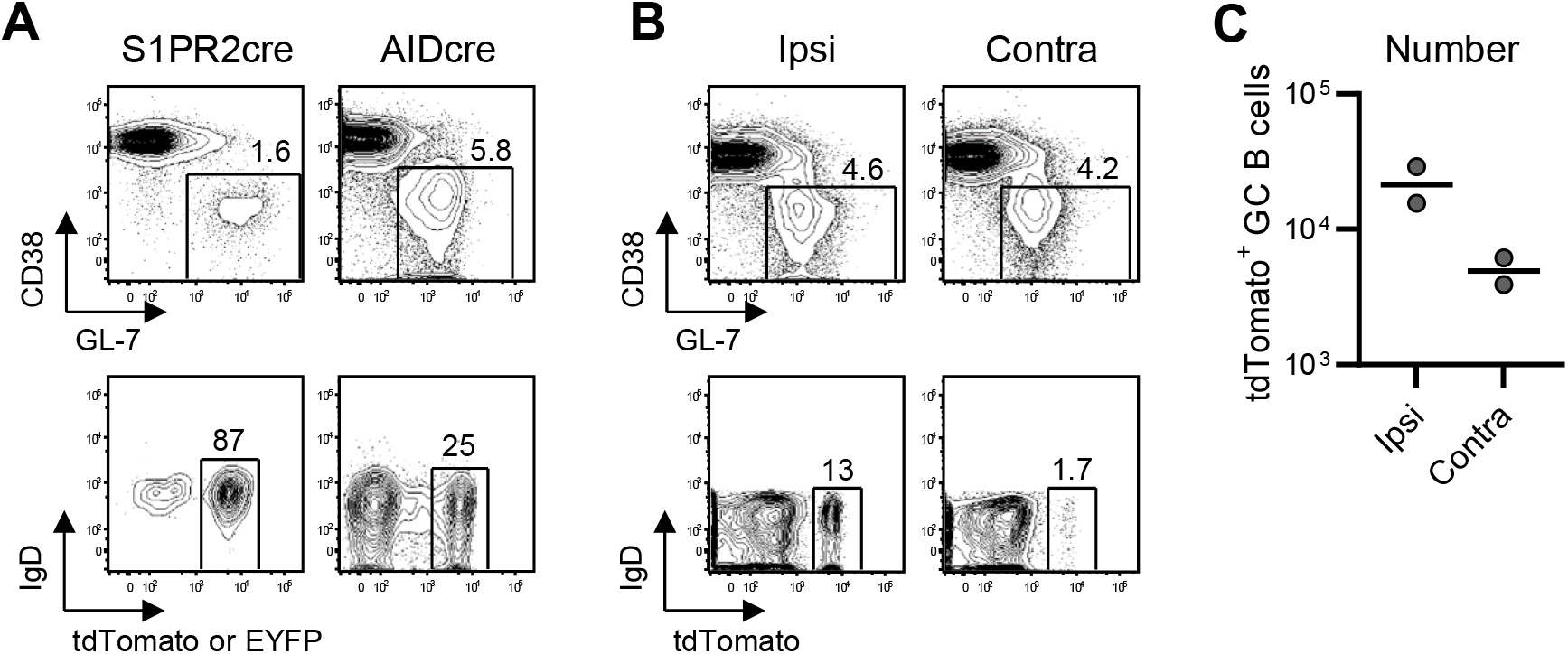
Tracing the fates of the progeny of primary GC B cells in *S1pr2*-ERT2cre-tdTomato mice. Participation of the progeny of primary GC B cells was assessed in fate-mapping mouse models. *S1pr2*-ERT2cre-tdTomato mice or AID-Cre-EYFP mice were primed with H1 SI-06, injected with tamoxifen (d8-d12). (**A**) Representative flow diagrams of GL-7 and CD38 expressions on B220+CD138-cells (top panels) and of tdTomato (left) or EYFP (right) and IgD expressions on B220+CD138-GL-7+CD38lo GC B cells (d14 primary, bottom panels) in *S1pr2*-ERT2cre-tdTomato mice (left) and in AID-Cre-EYFP mice (right). (**B** and **C**) Ten weeks after the priming, *S1pr2*-ERT2cre-tdTomato mice were boosted with homologous HAs (H1 SI-06) ipsilaterally or contralaterally. Mice were analyzed 8 days after boosts (see also figure legend for Fig. 4). Representative flow diagrams of GL-7 and CD38 expressions on B220+CD138-cells (top panels) and of tdTomato and IgD expressions on B220+CD138-GL-7+CD38lo GC B cells (**B**), and number of tdTomato+ secondary GC B cells following ipsilateral or contralateral boosts (**C**). Each dot represents an individual mouse. Horizontal bars, geometric mean.

**Figure S5.**
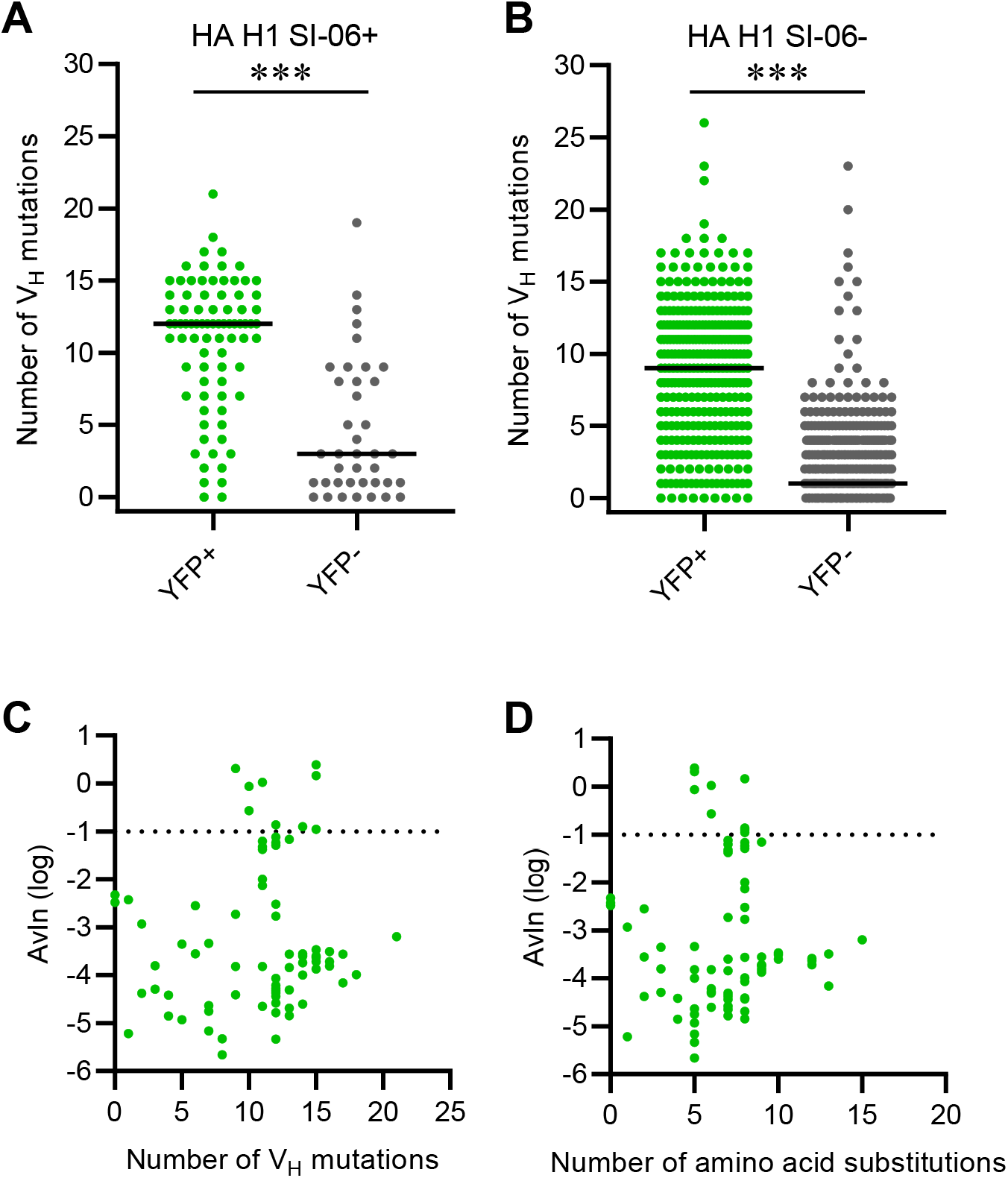
SHM of YFP+ secondary GC B cells in AID-Cre-EYFP mice. (**A** and **B**) Samples shown in Fig. 4D were split by the HA H1 SI-06 reactivity of respective clonal IgGs in culture supernatants. Distribution of the number of V_H_ point mutations recovered from HA-binding YFP+ (n = 82) and YFP-(n = 39) secondary GC B cells (**A**) and HA non-binding YFP+ (n = 317) and YFP-(n = 579) secondary GC B cells (**B**) following ipsilateral boosts. Horizontal bars represent mean. ***, *p* < 0.001 by Mann-Whitney’s U test. (**C**) Number of V_H_ mutations and AvIn values of respective clonal IgGs for HA-binding YFP+ secondary GC B cells (n = 82) were co-plotted. Horizontal dotted line represents AvIn = 0.1, which we consider high avidity. There are no significant correlation between number of V_H_ mutations and AvIn value (r = 0.14 and *p* = 0.22 by nonparametric Spearman correlation). (**D**) As in (**C**) with the number of amino acid substitutions. There are no significant correlation between number of V_H_ mutations and AvIn value (r = 0.18 and *p* = 0.09 by nonparametric Spearman correlation).

**Figure S6.**
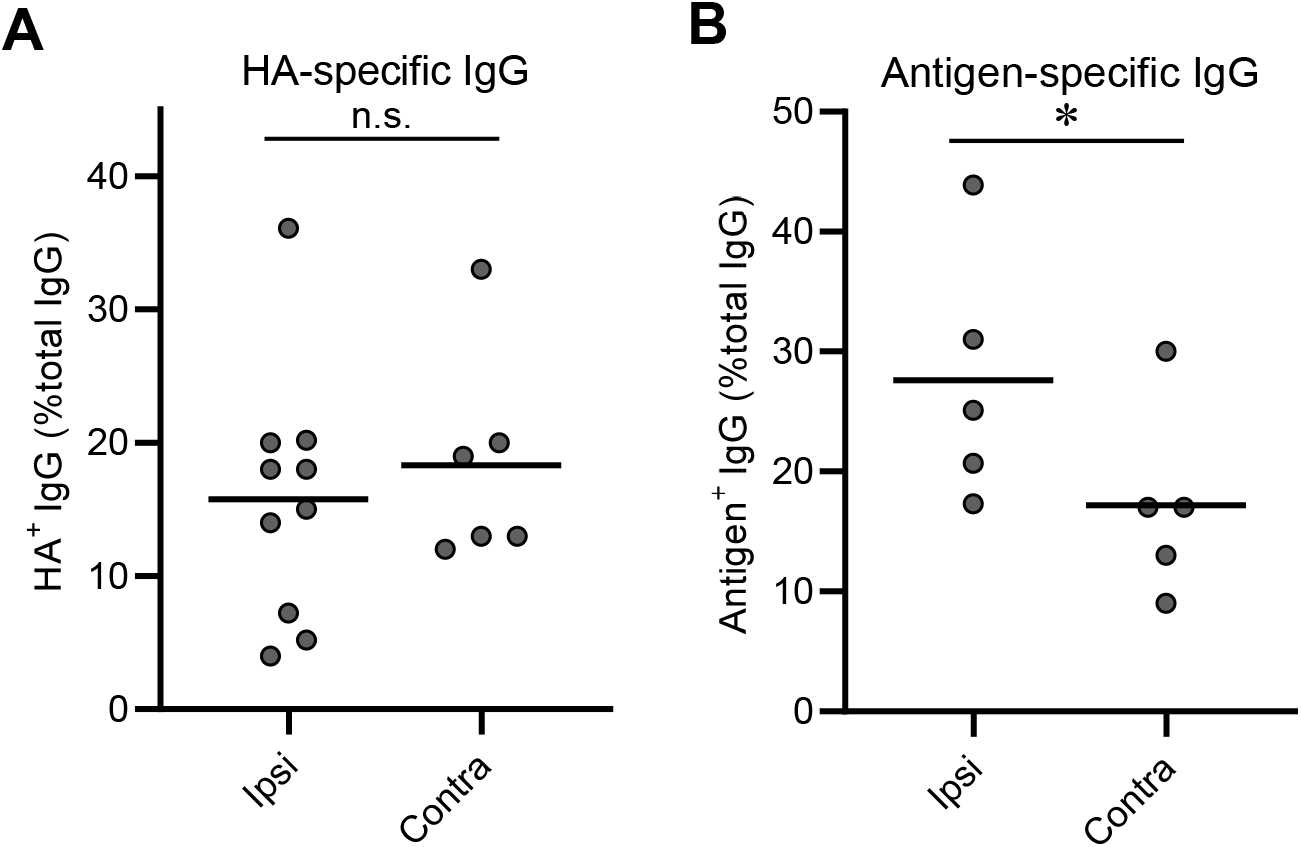
Frequency of antigen-specific IgGs among recall GC B cells. B6 mice were primed with H1 SI-06, and then 8-10 weeks later boosted with either H3 X31 (**A**) or rPA (**B**) ipsilaterally or contralaterally. Eight days following boosts, Nojima cultures were established for secondary GC B cells. After culture, HA-reactivity was determined for each clonal IgG by Luminex assay. Frequency of HA-specific IgGs among all clonal IgGs was calculated for each mouse sample, which is represented by each dot. *, *p* < 0.05; n.s., *p* > 0.05 by Mann-Whitney’s U test. Combined data from 4 (for **A**) and 2 (for **B**) independent experiments are shown.

**Table S1.**
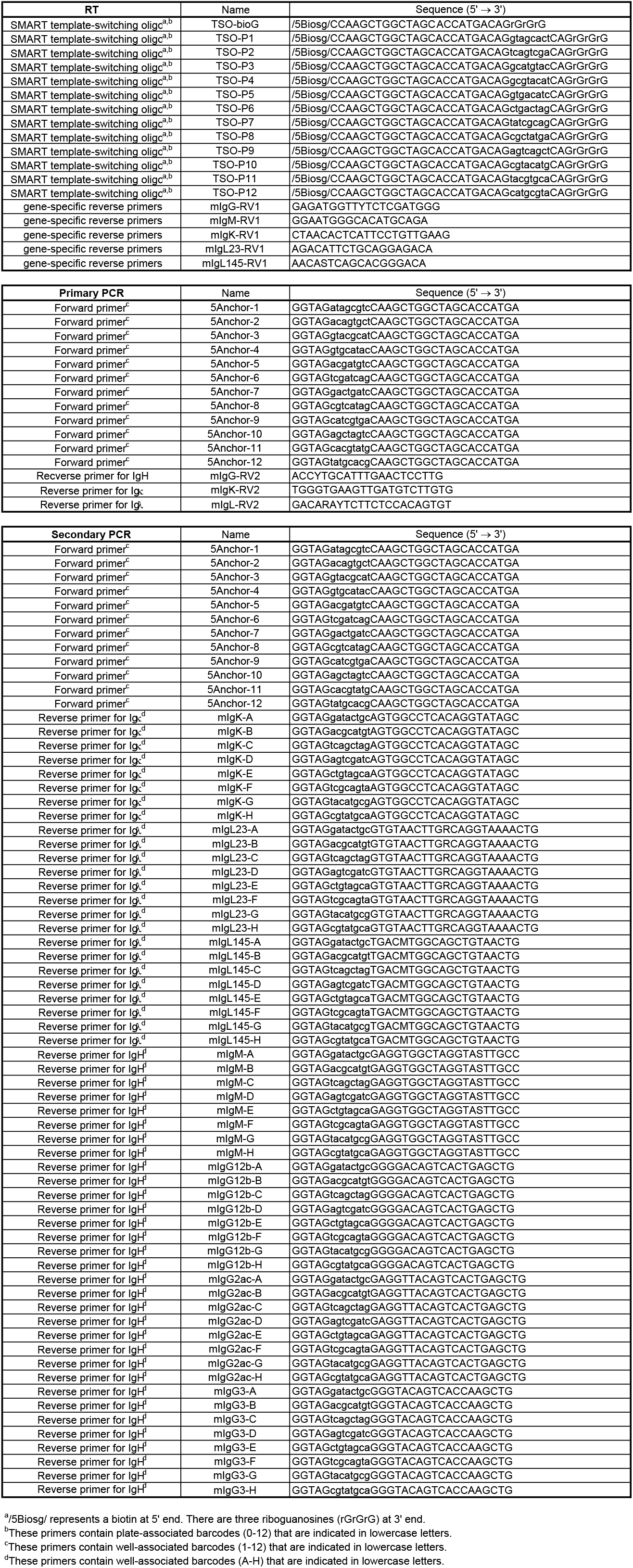
Primer sequences.

## Acknowledgments

We thank W. Zhang, X. Liang, D. Liao, A. Watanabe, and X. Nie (Duke University) for assistance. We thank Drs. C.A. Reynaud and J.C. Weill (Universite Paris-Descartes) for their gift of AID-Cre-EYFP mice. We thank Dr. T. Kurosaki (RIKEN) for *S1pr2*-ERT2cre-tdTomato mice. We thank Drs. R.W. Rountree, R. Spreng, and Y. Wang (Duke Human Vaccine Institutes) for advice on statistical analysis.

## Author contributions

M.K. and G.K. designed research. M.K., C-H.Y., and R.K. performed research. G.B. and S.C.H. provided reagents. S.S. and M.K. developed V(D)J sequencing method. M.K., C-H.Y., R.K., and G.K. analyzed data. M.K., S.C.H., and G.K. wrote the paper. All authors read and commented on the paper.

## Competing interests

Authors declare that they have no competing interests.

### Data and materials availability

All data are available in the main text or the supplementary materials.

